# Defining variation in pre-human ecosystems can guide conservation: An example from a Caribbean coral reef

**DOI:** 10.1101/749382

**Authors:** Aaron O’Dea, Mauro Lepore, Andrew H. Altieri, Melisa Chan, Jorge Manuel Morales-Saldaña, Nicte-Ha Muñoz, John M. Pandolfi, Marguerite A. Toscano, Jian-xin Zhao, Erin M. Dillon

## Abstract

There is a consensus that Caribbean coral reefs are a pale shadow of what they once were, yet a reef’s pre-human state is typically assumed or estimated using space-for-time substitution approaches. These approaches may fail to account for past variation before human impact which could mislead conservation priorities and actions. In this study we use a suite of fossilised mid-Holocene (7.2-5.6 ka) fringing reefs in Caribbean Panama to define the Historical Range of Variation (HRV) in coral community structure before human-impact to provide context for the states of modern reefs in the same area. Using the abundances of coral taxa to quantify communities, we found that most of the modern coral communities exist in novel ecosystem states with no fossil precedence. We do however identify one modern reef that is indistinguishable in coral community structure from the mid-Holocene reefs. Reef-matrix cores show that the community on this reef has remained in a stable state for over 760 years, suggesting long-term resistance to the region-wide shift to novel states. Without historical context this robust and stable reef would be overlooked since it does not fulfil expectations of what a “pristine” coral reef should look like. This example illustrates how defining past variation using the fossil record can place modern degradation in historical context and improve conservation recommendations.

## 1. Introduction

Caribbean coral reefs started to deteriorate long before most were first surveyed [1–7]. Consequently, modern benchmarks used to inform and evaluate the success of conservation actions often rely on space-for-time substitution, which assume that spatial patterns across modern gradients are a suitable proxy for temporal patterns. The approach is a mainstay of conservation ecology, and has proved beneficial in coral reef ecology [e.g. 8], but relies on the assumptions of uniformitarianism and that spatial and temporal variation are equivalent. Such assumptions have been challenged [9] and unlikely to be met in habitats that are naturally highly variable on small spatial and temporal scales [10]. Application of the space-for-time substitution approach to Caribbean coral reefs may be particularly problematic because (arguably) no pristine Caribbean reefs remain with which to establish true baselines. If true, conditions from one reef cannot be confidently used to define another’s baseline. Additionally, with few reconstructions of the state of reefs before human impact, baselines rely on limited definitions of reef states that likely fail to capture the high natural spatial and temporal complexity of coral reefs [3,11]. Accordingly, to prioritize conservation activities, contextualise restoration goals, and reveal the drivers of ecosystem change, we need to better quantify spatial and temporal variation, both past and present.

The Historical Range of Variation (HRV) is an approach developed in terrestrial ecology [12,13] to define past spatial and temporal variation in ecosystem states, thereby helping to determine if current states and rates of change fall within or outside the natural range of variation [14,15]. Coral reefs are particularly well-suited to the application of this approach given their high community complexity, and their propensity to preserve a wide array of reef community members [16–24] and past environmental signals [25] as they accrete. In this study, we quantify the HRV of coral communities in a suite of lagoonal fringing coral reefs in western Caribbean Panama (figures 1 & 2) using ∼7000-year-old (mid-Holocene) fossil assemblages. We then compare the HRV with the variation that exists on adjacent modern reefs today to quantify modern day ecosystem states within a historical context. Our findings provide a case study with which to discuss what should be considered natural and novel in ecosystems that are experiencing rapid change.

**Figure 1.**
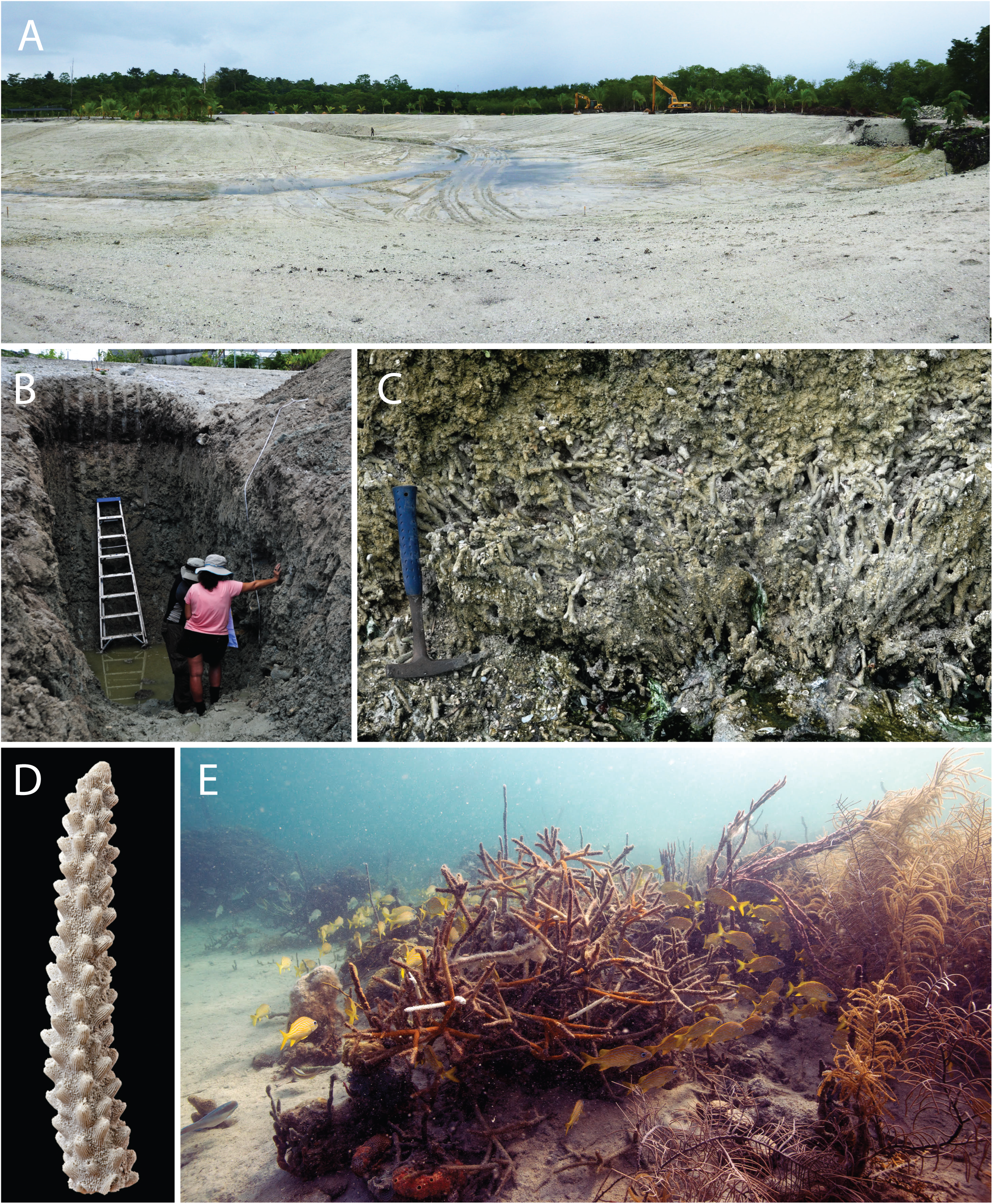
Fossil and modern coral reefs of eastern Almirante Bay, Caribbean Panama. (A) Large-scale excavation revealed the mid-Holocene fossil coral reef in Bocas del Toro, where we dug four trenches (B) to expose the autochthonous and in life position (C) fossil reef for bulk sampling. Preservation of corals, like *A. cervicornis* was excellent (D). The coral community composition at the modern reef at Punta Caracol (E) was found to be encapsulated by the variation described in the mid-Holocene reefs. Images courtesy of Harry Taylor (D) and David Kline (E).

**Figure 2.**
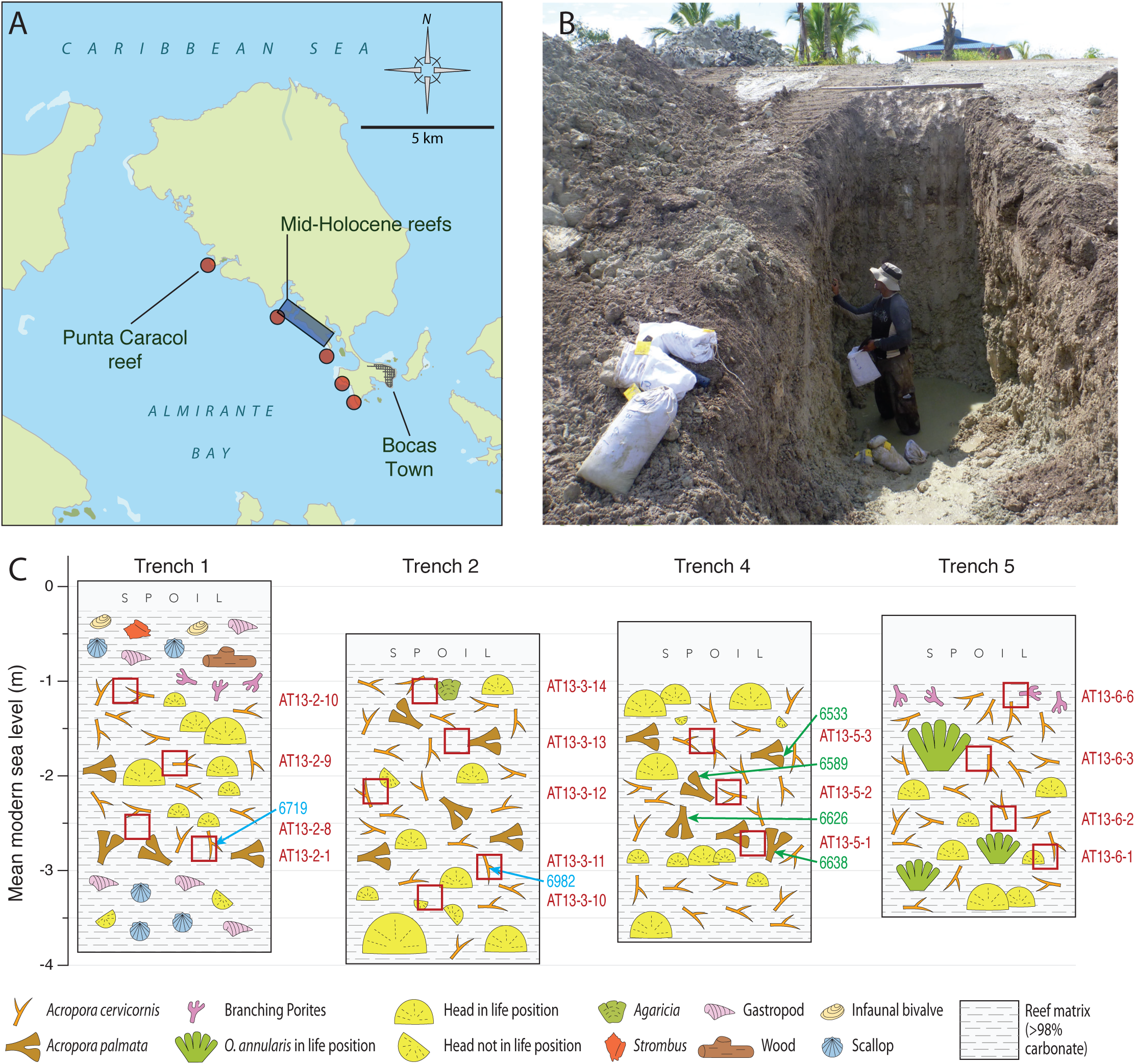
(A) Map of study area showing the location of modern reefs sampled (red circles) and known extent of mid-Holocene reef (blue rectangle). Map is centered on 9.359395 N, 82.319180 W. (B) Example of the Trenches (#4) in the mid-Holocene reef made to permit bulk sampling of *in situ* reef framework. (C) Stratigraphic sections of the trenches showing dominant taxa and positions of bulk samples (red squares), U-Th dates (blue) and radiocarbon dates (green), both years BP.

## 2. Materials and Methods

### 2.1. General setting

Fringing coral reefs, seagrass and mangrove systems became established in the Almirante Bay (figure 2*a*) shortly after the bay flooded around 8000 years ago when sea levels rose [18] and have persisted since [1,26,27]. Today, the reefs fringing the bay are primarily composed of branching *Porites* and *Agaricia* with moderate to low coral cover and fish communities that are depauperate of adults [28]. Almirante Bay is well protected from wave action and currents and sits outside the hurricane belt. High rainfall, deforestation and changing land use, including a ∼140-year banana industry [29], cause relatively higher suspended particles and dissolved nutrients conditions compared to other Caribbean reef systems [30,31]. The bay today experiences periodic hypoxia that sometimes causes widespread death of corals and other aerobic reef-inhabiting organisms [32,33].

### 2.2. The fossil reef

Two large-scale developments led to the excavation of ∼50 hectares of a fossil fringing coral reef in the eastern portion of Almirante Bay (figures 1 & 2). Previous Uranium-Thorium (U-Th) dates from coral pieces placed the fossil reef in the mid-Holocene, ∼7 ka [18]. Humans were on the Isthmus of Panama before 14 ka [34,35] but remained concentrated on the Pacific coast until major settlements of people exploiting Caribbean marine resources appeared after 4 ka [36–38]. As such, this large fossil coral reef offers an opportunity to quantitatively reconstruct the spatial and temporal variation in community structure in a fringing reef system from a time before major human impact.

### 2.3. Quantifying the structure of mid-Holocene and modern coral communities

We mapped ∼11 hectares of the known extent of the mid-Holocene reef site using high-resolution Trimble R8 Base and Rover GNSS system to build a 3D model of the site (called Sweet Bocas) and locate major fossil habitats (see electronic supplementary material, figure S1, for details). Using a large excavator, we dug four 4-5 m deep and ∼1.5 m-wide trenches into the reef to expose fresh *in situ* reef framework (figures 1 & 2; electronic supplementary material, figure S2). Sections were measured and mapped and the vertical positions of both bulk samples and coral samples for dating were measured using the Trimble R8 system (electronic supplementary material, figure S1). All elevations were adjusted to mean sea level (MSL) by measuring sea level directly adjacent to the fossil site and relating that to data from the tide gauge at the Smithsonian Tropical Research Institute’s Bocas Research Station.

To quantify the HRV in coral communities in the mid-Holocene fossil reefs, we extracted 16 bulk samples (∼10 kg each) of reef framework from each of the four trenches we excavated (figures 1b, 2b). To compare the resulting HRV with modern reef communities, we collected 22 bulk samples (∼10 kg each) of reef framework from the top < 10 cm of the death assemblage from five modern fringing reefs adjacent to the fossil site (figure 1e). These modern death assemblage samples were collected at water depths between 2 and 10 m to ensure we covered the full range of depths that we sampled in the mid-Holocene. Depth of reef formation in the mid-Holocene was almost certainly within this range because of the presence of *Acropora palmata* in most sections (electronic supplementary material, figure S3).

In both mid-Holocene and modern bulk samples, all coral skeletal remains in the > 2mm fraction were identified to species or genus using the Caribbean Coral Skeleton Identification Guide [39] – a purpose-built reference collection – and other guides [40]. The relative mass of each coral taxon was square-root transformed and used to compare the assemblage structure of each sample using Bray-Curtis dissimilarities, which were ordinated using non-metric multidimensional scaling (NMDS) to place samples in multivariate ecological space based on the relative abundance of coral taxa skeletal weights [39]. *PERMANOVA* was used to test for significant differences in dissimilarities among groups. Corals of unknown taxa were removed from all analyses. These corals are listed as “unknown” in electronic supplementary material, Table S2, and were generally fragments identified as coral but too eroded or fragmented to be able to be confidently placed in a taxon [39]. “Unknown” corals totalled 8.26% of all coral weights in mid-Holocene samples and 10.02% in modern samples (electronic supplementary material, Table S2), suggesting no substantial difference in preservation between the two ages. Relative weights of coral taxa in each sample were square-root transformed to reduce the influence of dominant taxa prior to analysis. To calculate Bray-Curtis dissimilarities of samples we used the function *vegdist* of the software package *vegan* [41] in *R*. The NMDS (Kruskal’s Non-metric Multidimensional Scaling) was computed using *isoMDS* function of the software package *MASS* [42] in *R* which was then plotted using *ggplot* [43] in R [44]. The function *adonis* (Permutational Multivariate Analysis of Variance Using Distance Matrices) of the software package *vegan* [41] was used to test for significant differences in dissimilarities among groups. To confirm that differences between communities were valid, and not an artefact of differences between groups in how much each community deviates from the centroid of the group we used beta-dispersion with the *betadisper* function [Multivariate homogeneity of groups dispersions (variances)] in the software package *vegan* [41]. All analyses were performed in *R* (R core team).

### 2.5. Chronological framework

We used radiometric dating (U-Th and radiocarbon) to provide temporal context to our data. To determine how old and for how long the mid-Holocene reef grew, we used radiocarbon dating on eight pieces of *Acropora palmata* and U-Th on two branches of *A. cervicornis* (electronic supplementary material, Table S1) and combined these results with nine U-Th dates previously conducted on large coral heads from the same site [18]. To estimate how much time is encapsulated in a modern bulk sample we used U-Th dating on six pieces of *A. cervicornis* randomly selected from one of the bulk samples from the modern reef (i.e. a death-assemblage) at Punta Caracol reef. See electronic supplementary material for further information on radiometric dating methods.

## 3. Results and Discussion

### 3.1. Preservation, age and time-averaging

Corals from the mid-Holocene fringing reefs were found to be, on the whole, very well-preserved with chemically pristine skeletal material [18] (figure 1d). The proportion of colonies eroded to the point that they were unidentifiable, a proxy for taphonomic preservation, was slightly higher in the modern reefs (10.02%) than the fossil reefs (8.26%) (electronic supplementary material). The majority of fossil colonies were found preserved in life-position in unsorted fine carbonate muds and silts (figures 1 & 2; electronic supplementary material figure S2), suggesting that these fossil reefs comprise autochthonous assemblages that accumulated without major disturbance. These mid-Holocene reefs are located inside the bay on the leeward side of >20 m high Pliocene bedrock rise, suggesting that the setting that they grew in was similarly a highly protected lagoon that the modern reefs occupy today. U-Th and radiocarbon dating reveal that the sections of the reefs that we sampled grew, without apparent interruption, over a period of at least 1,600 years (7.2-5.6 ka) (electronic supplementary material Table S1), when sea-level had reached similar levels to today.

U-Th dates from a bulk sample taken from a modern reef with living coral ranged in age from 1926 to 2012 AD (mean = 1991 AD, SD = 30.4 years, n = 6; electronic supplementary material, Table S1), suggesting that when they are actively accreting, coral assemblages in reefs with branching framework are predominantly formed by modern corals (i.e., post 1950). This finding supports previous work showing little vertical mixing in reef matrix comprised of branching corals that accumulate in protected areas [20,21,45], presumably because the branching coral limits infaunal and epifaunal bioturbation which can cause considerable vertical mixing in other settings [46,47].

### 3.2. Coral community composition in the mid-Holocene and today

We used the relative abundances (weights) of coral taxa in the fossil samples to define the HRV of the reefs in the mid-Holocene and compare that HRV with modern community composition on the adjacent reefs today. Ordination of the data demonstrates that, overall, the modern coral communities in Almirante Bay are distinct from the fossil defined HRV (figure 3). The ecological space occupied by mid-Holocene coral communities, estimated by NMDS, is significantly different (PERMANOVA; p <0.0001) from modern coral communities in the same area. Coral communities in this reef system today are therefore distinct from their mid-Holocene counterparts, suggesting that the modern reefs exist in novel states.

**Figure 3.**
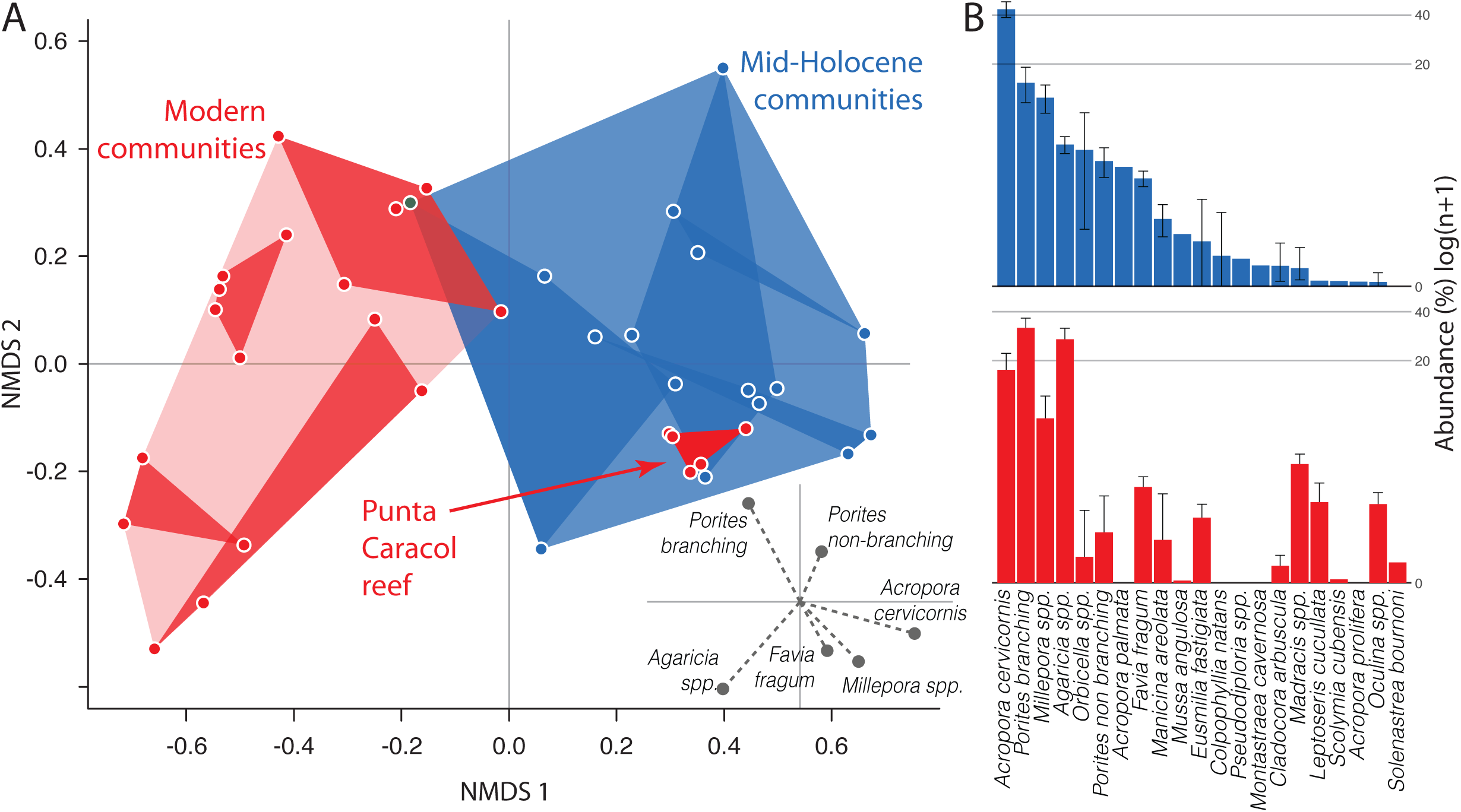
Most modern coral communities (red) are compositionally distinct from their fossil counterparts (blue) except the living reef at Punta Caracol. (A) Non-metric multidimensional scaling (NMDS) ordination place communities in multivariate ecological space based on the relative abundance of coral taxa skeletal weights [39]. Inset shows the vector and magnitude of the scores from the six most explanatory taxa. (B) Rank abundance plots (log-scale) show shifts in the dominant and rare taxa between mid-Holocene (top) and modern (bottom) reefs.

Mid-Holocene reefs were generally dominated by staghorn coral (*Acropora cervicornis*), which was rare or absent on many of the modern reefs (electronic supplementary material, figure S4). This finding is consistent with other paleoecological work showing that acroporid corals were the principal pre-Anthropocene reef-builders in the Caribbean [48,49] before the historical [1] and more recent [23] declines. Modern coral communities on the fringing reef we studied are instead primarily dominated by finger coral (branching *Porites* spp.) or lettuce leaf coral (*Agaricia* spp.) (figure 3), corroborating results from benthic surveys that show *Agaricia* and branching *Porites* to have dominated these reefs, and others across Almirante Bay, for at least the last 20 years [30,50,51]. This trend mirrors the Caribbean-wide transition from acroporid to *Agaricia* and branching *Porites* dominance [51–53] frequently attributed to *Agaricia* and *Porites* being more robust to stress and disease than *Acropora.*

### 3.3. A resilient reef at Punta Caracol

In contrast to the shift to novel ecosystem states observed in all of the other modern reefs sampled in this study, we found that the coral community composition at one modern reef – Punta Caracol – is entirely contained within the fossil-defined HRV (figure 3). Like the mid-Holocene reefs, coral assemblages at Punta Caracol are dominated by *A. cervicornis* (electronic supplementary material, figure S4). Benthic surveys show that *A. cervicornis* has existed on the reef since at least 1997 [50,54]. To reveal the deeper history of this reef, we collected two push-cores following methods described in Cramer et al. [20]. Cores were taken ∼2 m apart from each other at a water depth of 2.2 m on top of living *Acropora cervicornis* stands. 5 cm-deep samples were extracted from the core matrix every 10 cm, sieved to >2 mm and all coral pieces identified [39]. Results show over 760 years of dominance by *A. cervicornis* without apparent interruption (electronic supplementary material, figure S5, table S1). When taken together, these findings suggest that Punta Caracol is a reef that has evaded the shift to a novel state that has occurred on other reefs in the area.

We do not claim that the reef at Punta Caracol is pristine. It is unlikely that the reef could have avoided the impacts of conch [55], lobster and fish harvesting [37], regional and global pollution [30] and global change in water temperature and chemistry [32]. Nevertheless, in spite of these stressors, the coral community has evaded the region-wide shift to a novel state that has impacted other reefs in the area. Determining why Punta Caracol exists in the state it does today could reveal information that can be repurposed for the conservation of other reefs. Surprisingly though, we see no clear differences in the reef’s abiotic characteristics (i.e. topography, water flow, water residence time, frequency of hypoxia, suspended solids, dissolved nutrients and temperature) or biotic characteristics (i.e., planktonic productivity, coral cover, fish abundance) [30,32,56,57] when compared to other reefs in the region that could account for why Punta Caracol is exceptional. Measuring local anthropogenic stressors such as fishing pressure, pollution and tourism was outside the scope of this paper, but of all the modern reefs studied here the Punta Caracol reef is located farthest from the densest human population in the region (figure 2), implicating a lack of local-human disturbance as the reason for the reef’s resilience. Alternatively, Punta Caracol could be a fortunate ‘oasis’ [58], although the 760-year stability in community composition (electronic supplementary material, figure S5) undermines such a stochastic explanation, which would likely have manifested in shifting coral community composition over time. Finally, feedback mechanisms, such as the positive density-dependent increase in resilience of dominant patches of *A. cervicornis [59]* (up to certain densities [60]) might have played a role in maintaining community stability on the reef. *A. cervicornis*, like most acroporid corals, is a species that has traded disease and predation-resistance for fast-growth and high-rates of clonal reproduction [23]. Previous work has shown that the population of *A. cervicornis* at the Punta Caracol reef has particularly low genetic diversity relative to other conspecific populations in Bocas del Toro region, a sign that it has been isolated from other populations and established principally through clonal propagation [54]. Could a resilient, genetically isolated population of *A. cervicornis* have persisted for more than 760 years at Punta Caracol through clonal propagation? Historical information from Punta Caracol could be used to make more informed choices when selecting resilient genets to translocate coral colonies, improve connectivity and recruitment [61] and promote recovery [62].

### 3.4. Past variation as a guide to future conservation

There is a growing trend in conservation biology to accept that the majority of ecosystems today have already shifted into irreversible novel states and, consequently, that conservation resources should not be squandered trying to coerce restoration to unachievable historical conditions. For example, the near-ecological extinction of megafauna and fish from overharvesting and the chronic and intensifying effects of successive coral bleaching make returning Caribbean coral reefs to their past states currently unimaginable. While in general this is clearly true, this paradigm involves the assumption that historical conditions of Caribbean coral communities were universally of high coral cover and high fish biomass [63], a notion that persists even though reefs exhibit considerable variation in these measures today [2] and did so in the past [16]. This assumption is reinforced by space-for-time substitution studies that take reefs that are perceived to be healthy and presume their states are the baselines for reefs perceived to be degraded. Our study highlights the pitfalls of the use of space-for-time substitution approaches in coral reefs. Not even an experienced reef ecologist is likely to think that Punta Caracol resembles a pristine Caribbean reef. Living coral cover and fish biomass, often used as a comparative metric of reef health, are not even the highest observed on the reefs in the bay [30]. Yet despite failing to fulfil expectations of what a “pristine” coral reef should look like, we demonstrate, through the use of the HRV approach, that the coral community at Punta Caracol has persisted for centuries in a state that retains at least some elements of their prior condition before human impact.

Defining the HRV of an ecosystem using the fossil record can therefore provide powerful, location-specific context for modern ecosystem change (figure 4), which can provide clues as to the ecosystem types, localities and genets that could confer functional, taxonomic and genetic elements useful for conservation. This could be valuable because it is difficult to predict how complex ecosystems like coral reefs will change in the future because of processes like extinction debt [64], the unclear directionality of future ecosystem shifts [65] and the complex impact of unidentified future stressors [66]. Conservationists interested in supporting coral reefs must therefore hedge their bets by conserving variability itself to provide nature with the raw material for the selection of resilience [67]. Resources for conservation will always be limited and decisions must be made about where priority should be given. We suggest that such decisions be made not only based on current states and future predictions, but also through the use of HRV data to extend time scales, establish location-specific baselines and uncover elements of resilience that might otherwise be overlooked.

**Figure 4.**
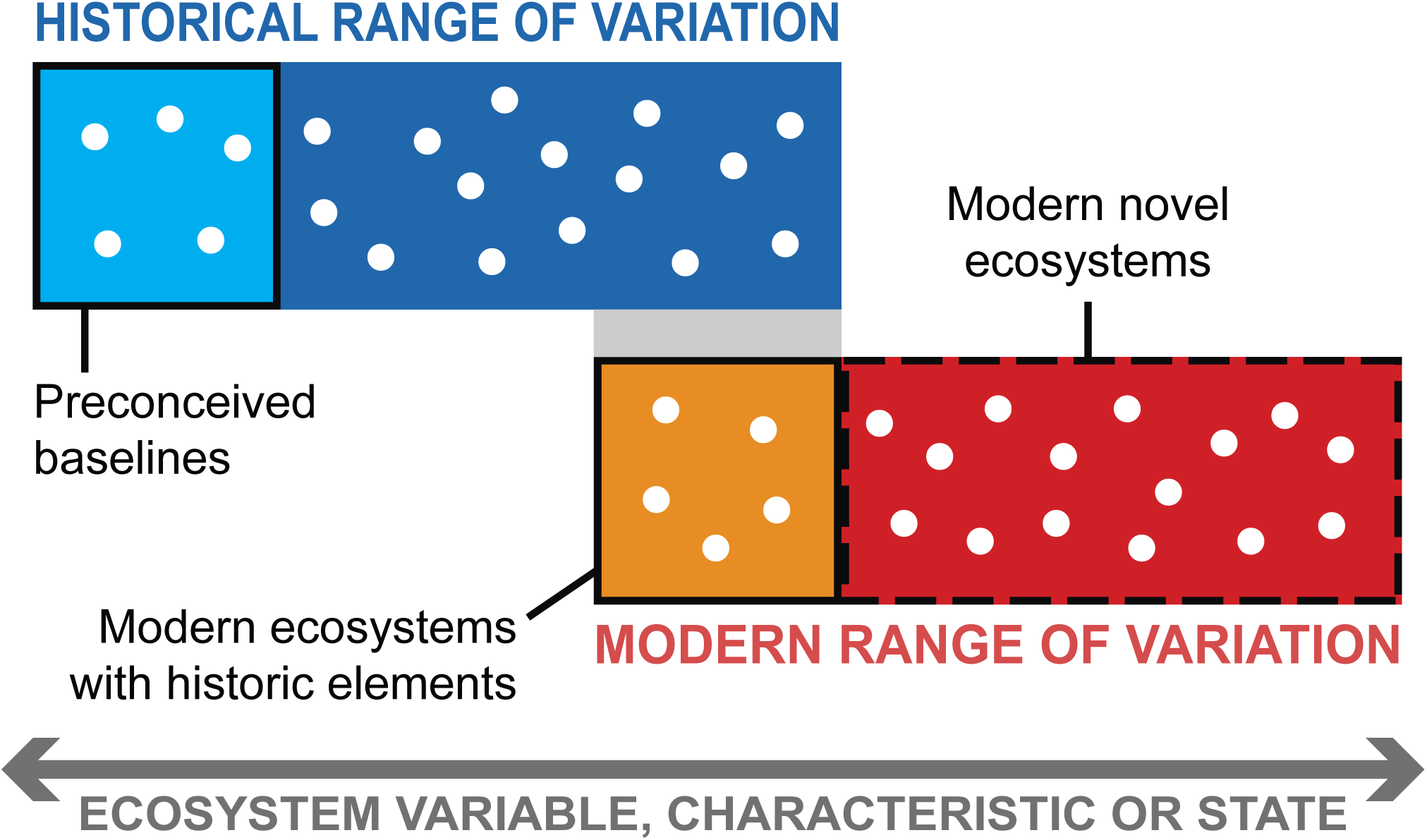
Quantifying the Historical Range of Variation (HRV) using fossils and comparing them to modern ecosystems can redefine preconceptions of pre-human ecosystems, establish which ecosystem states are truly novel, and reveal ecosystems that retain elements of their prior condition before human impact. Cartoon of localities (dots) along a gradient of ecosystem state, characteristic or other measurable ecological or environmental variable.

## Data accessibility

All data supporting the findings of this study are available within the paper and its electronic supplementary material files.

## Authors’ contributions

AO and ML conceived the study; AO, JM, MT and ED Collected material; AO, ML, AA, MC, JM, and N-H.M. identified material; AO. ML, MC, JM, N-HM, MT and J-xZ collected data; AO. ML, MC, N-HM, MT and J-xZ developed and implemented analyses, AO, ML, AA, MC, JM, N-HM, JP, MT, J-xZ and ED wrote and edited the manuscript.

## Acknowledgements

A. Belanger, A. Villarreal and the staff at Sweet Bocas kindly gave access to the fossil site and provided much logistical support for which we are grateful. For their many contributions, we thank U. Gonzalez, P. Gondola, G. Jacome, the Bocas Research Station Staff, J. Mate, C. De Leon, S. Dos Santos, F. Rodriguez, B. de Gracia, E. Grossman, S. Finnegan, D. Doughty, M. Alvarez, F Alvarez, D. Kline, S. Vollmer, J. Edlinger, M. Hynes, X. Bella Medrano, N. Downer, S. Mattson, G. Cumming, B. Figuerola, A. Endara, R. Solis, M. Leray, S. Finnegan, D. Norris, H. Guzman, N. Knowlton, M. Pierotti, G. Roff, N. Leonard, J.B.C. Jackson and K. Cramer. Fieldwork and U/Th dating was supported by donations from M. Selin and family, J. Bilyk, V. and B. Anders, J. and M. Bytnar and the Young Presidents Organization (Los Angeles Gold Chapter).

