## Supplemental information for "Defining variation in pre-human ecosystems can guide conservation: An example from a Caribbean coral reef"

**Electronic Supplementary Material for**

**This file includes**

Supplemental radiometric dating methods

figures S1 to S4

Tables S1 and S2

References for supplementary material

**Supplemental radiometric dating methods**

U–Th dating was conducted on coral fragments using a Nu Plasma multi-collector inductively-coupled plasma mass spectrometer (MC-ICP-MS) in the Radiogenic Isotope Facility at the University of Queensland, following chemical treatment procedures and MC-ICP-MS analytical protocols described in Clark et al. [1]. For U–Th dating, each coral sample consisting of ~150 mg fine sand-size chips that were carefully fragmented, $H_2O_2$-treated and then hand-picked under a binocular microscope to remove any trace detritus or grains with discolouration was spiked with a mixed $^{229}Th$–$^{233}U$ tracer and then completely dissolved in double-distilled concentrated $HNO_3$. After digestion, each sample was further treated with $H_2O_2$ to decompose trace amounts of organic matter (if any) and to facilitate complete sample-tracer homogenization. U and Th were separated using conventional anion-exchange column chemistry using Bio-Rad AG 1-X8 resin. After stripping off the matrix from the column using double-distilled 7N $HNO_3$ as eluent, $_4$N $HNO_3$ and 2%$HNO_3$ +0.03% HF mixture were used to elute U and Th into 3.5 ml pre-cleaned test tubes, respectively. After screening U and Th concentrations in the U and Th fractions using their 1:100 dilute solutions on a quadrupole ICP-MS, the U and Th separates for the samples were then re-mixed in 2% $HNO_3$ to make ~3 ml solution in a pre-cleaned 3.5 ml test tube. The mixed solution for each sample contained the entire Th fraction and a small percentage of the U fraction. The amount of U fraction to be added to the mixed solution was calculated based on the screening results and the MC-ICP-MS working sensitivity of the day, aiming to achieve ~5 volts of $^{238}U$ signal. The remixed U–Th solution was injected into the MC-ICP-MS through a DSN-100 desolvation nebulizer system with an uptake rate of ~0.06 ml/min. U–Th isotopic measurements were performed on the MC-ICP-MS using a detector configuration to allow simultaneous measurements of both U and Th isotopes. The $^{230}Th/^{238}U$ and $^{234}U/^{238}U$ activity ratios of the samples were calculated using the decay constants given in Cheng et al. [2]. $^{230}Th$ ages were calculated using the Isoplot 3.75 Program [3]. $^{230}Th$ ages were corrected for non-radiogenic $^{230}Th$ contributions using a modelled two-component-mixing non-radiogenic $^{230}Th/^{232}Th$ value based on the equation in Clark et al. [1].

We also obtained four AMS $^{14}C$ dates on exceptionally-well preserved branches of Acropora palmata (electronic supplementary material figure S4) from Trench 4 (figure 2). A. palmata generally occupies a narrow range of <1 to 5 (but can be up to 15) m water depth [4,5], but in the low-visibility waters of Almirante bay has only been observed in depths <2 m (A. Castillo, pers. comm.). Robust branches were found in the trenches at elevations of -3 to -1.5 m below mean modern sea level in some of the trenches (figure 2). Samples for dating were washed and air-dried, cut into slabs, and subsampled to retrieve pristine interior material without encrusters, epibionts and submarine cements. Prior to dating, X-Ray Diffraction was undertaken to determine if the samples consisted of 100 percent Aragonite; all samples were chemically pristine with no recrystallization or internal cements. Cleaned coral pieces were radiocarbon-dated at University of Georgia Center for Applied Isotope Studies. Coral pieces were first treated with dilute HCl to remove any contamination from the surface. The washed and dried sample was treated in the vacuum with concentrated phosphoric acid to recover carbon dioxide. For accelerator mass spectrometry analysis the cleaned samples were combusted at 900°C in evacuated / sealed ampoules in the presence of CuO. Resulting carbon dioxide was cryogenically purified from the other reaction products and catalytically converted to graphite using the method of Vogel et al. [6]. Graphite $^{14}C/^{13}C$ ratios were measured using the CAIS 0.5 MeV accelerator mass spectrometer. The sample ratios were compared to the ratio measured from the Oxalic Acid I (NBS SRM 4990). The sample $^{13}C/^{12}C$ ratios were measured separately using a stable isotope ratio mass spectrometer and expressed as $\delta^{13}C$ with respect to PDB, with an error of less than 0.1‰. The quoted uncalibrated dates have been given in radiocarbon years before 1950 (years BP), using the $^{14}C$ half-life of 5568 years, with error quoted as one standard deviation reflecting both statistical and experimental errors. The date has been corrected for isotope fractionation. Recovered dates were calibrated to calendar years before 1950 (cal BP) using CALIB 7.02 [7,8], the Marine13 calibration dataset [9], and an integrated, time-dependent global ocean reservoir correction. The age difference between the local ocean reservoir and modeled values (DR) was set at −36 ± 35 yr for the Gulf of Mexico [10]. One and two sigma calibrated ranges are reported with CALIB median probability values (electronic supplementary material, Table S1).

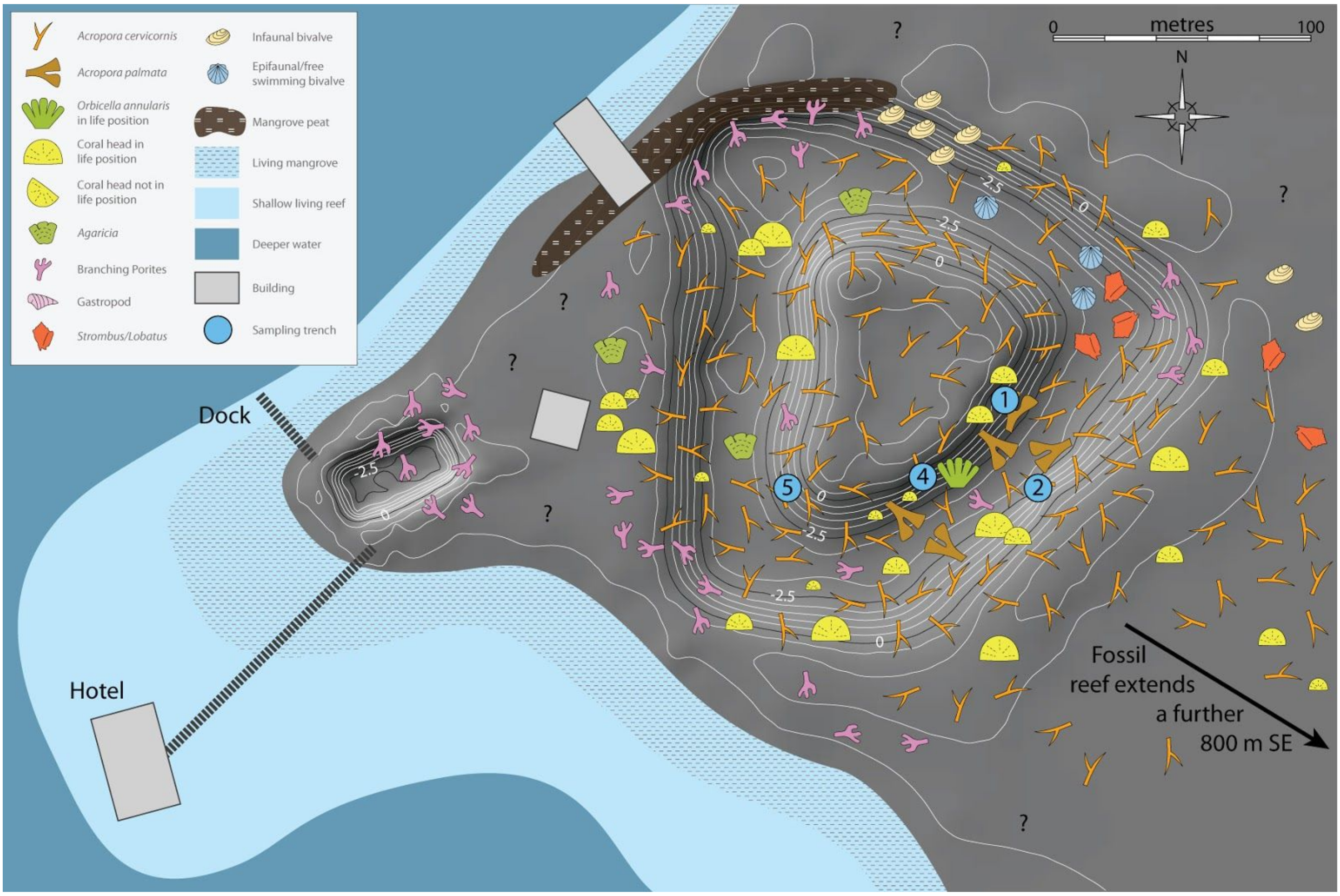


Figure S1. 3D model and habitat (facies) map of the NW portion of the mid-Holocene fossil reef site at Sweet Bocas, Bocas del Toro, Panama. Distribution of the remaining parts of the fossil reef are detailed in [11]. Contours in metres relative to MSL (see Materials and Methods for further information). The location of dominant fossils in the exposed surface matrix are labelled and the location of excavated trenches used in this study are marked. Map is centred on 9.359898°, -82.272677°.

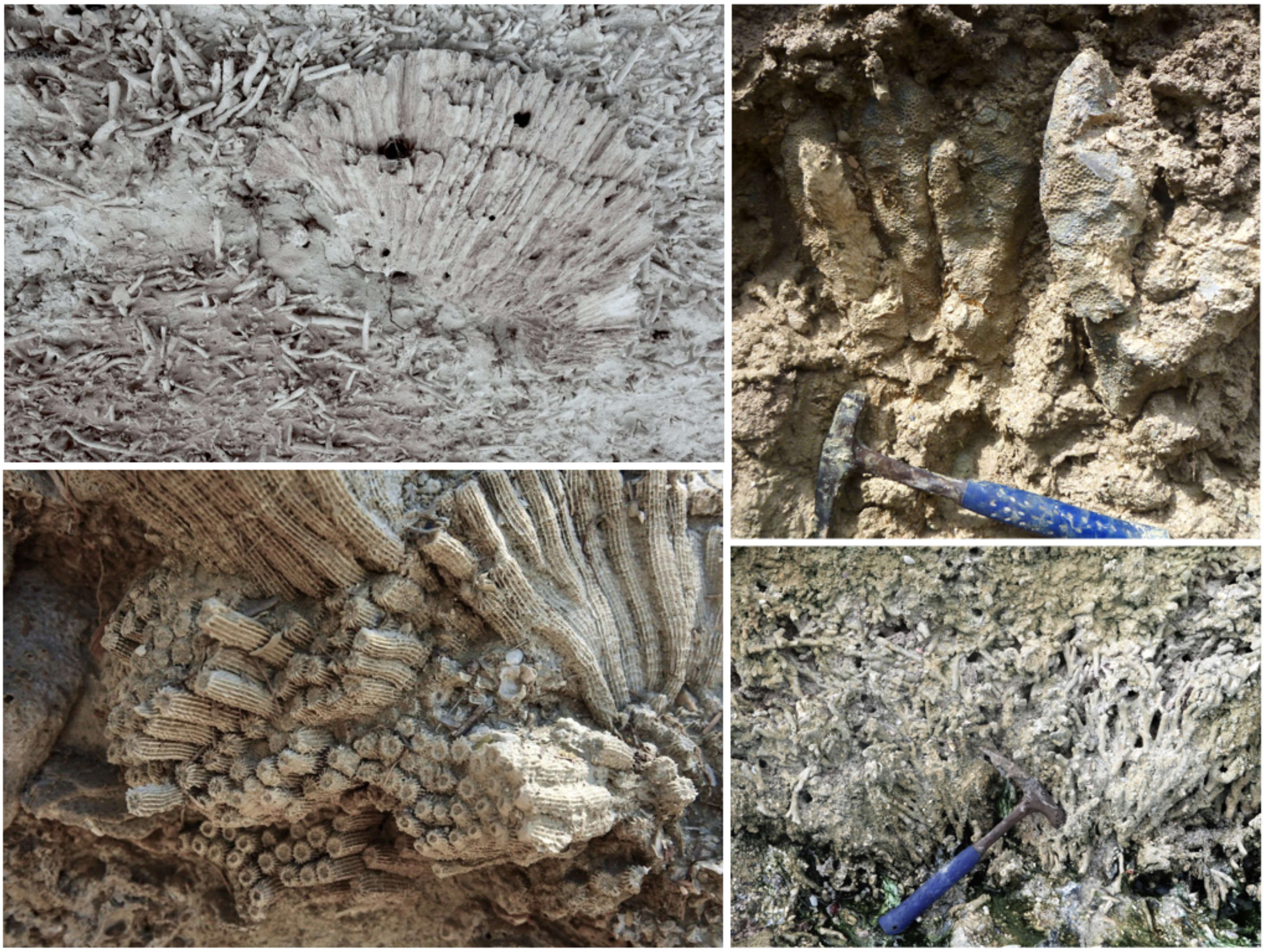

Figure S2. *In situ* and in life position corals exposed after trenching the mid-Holocene reef at Sweet Bocas, Bocas del Toro, Panamá**.** (a) a large coral head (possibly *Diploria*) in life position surrounded by fragments of *Acropora cervicornis*. (b) Columnar *Orbicella annularis* colony in life position. (c) Colony of *Montastraea cavernosa* in life position. (d) Large colony of branching *Porites sp.* in life position.

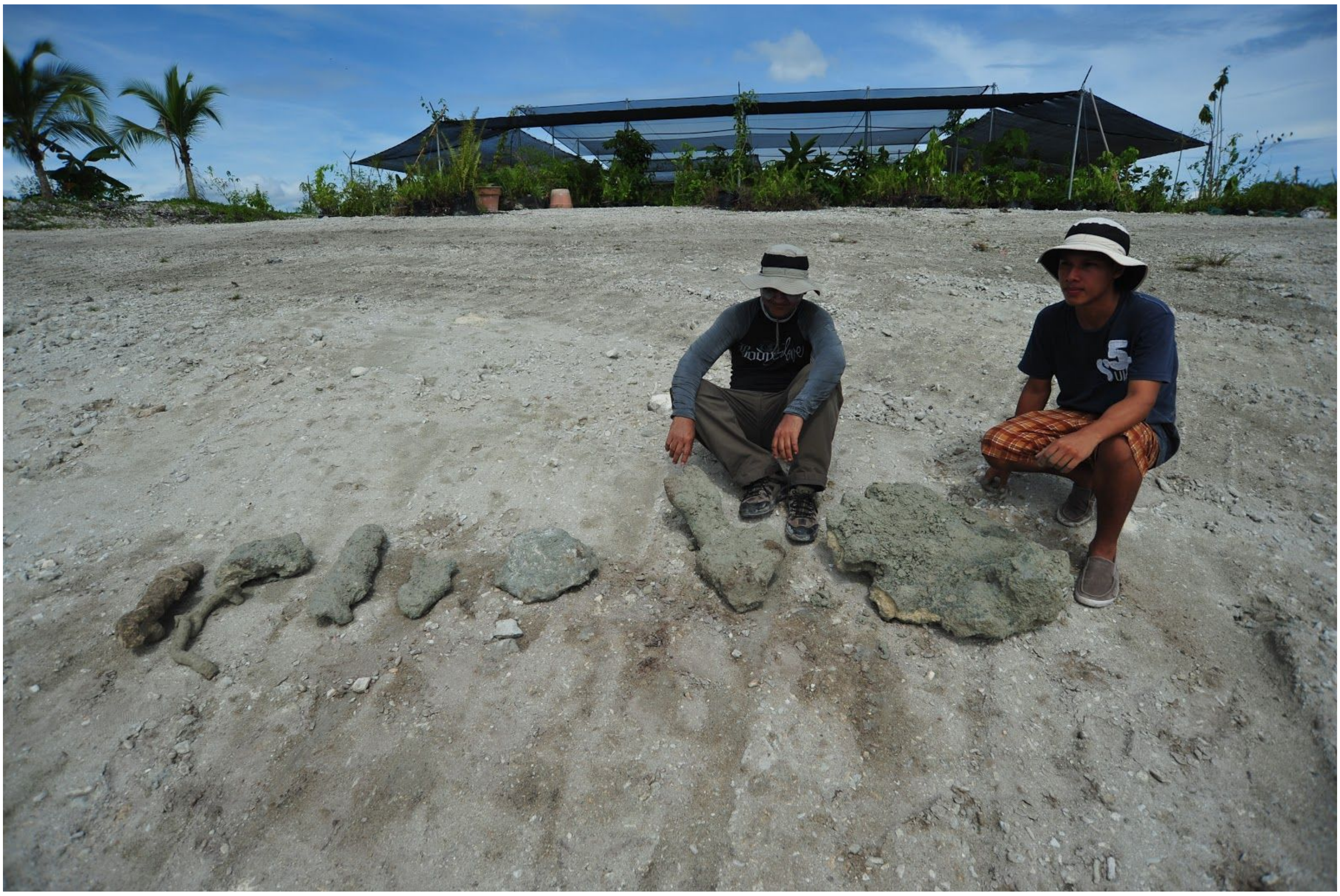

Figure S3. Examples of the many large fronds of well-preserved, chemically-pristine *Acropora palmata* colonies found in the mid-Holocene reef.

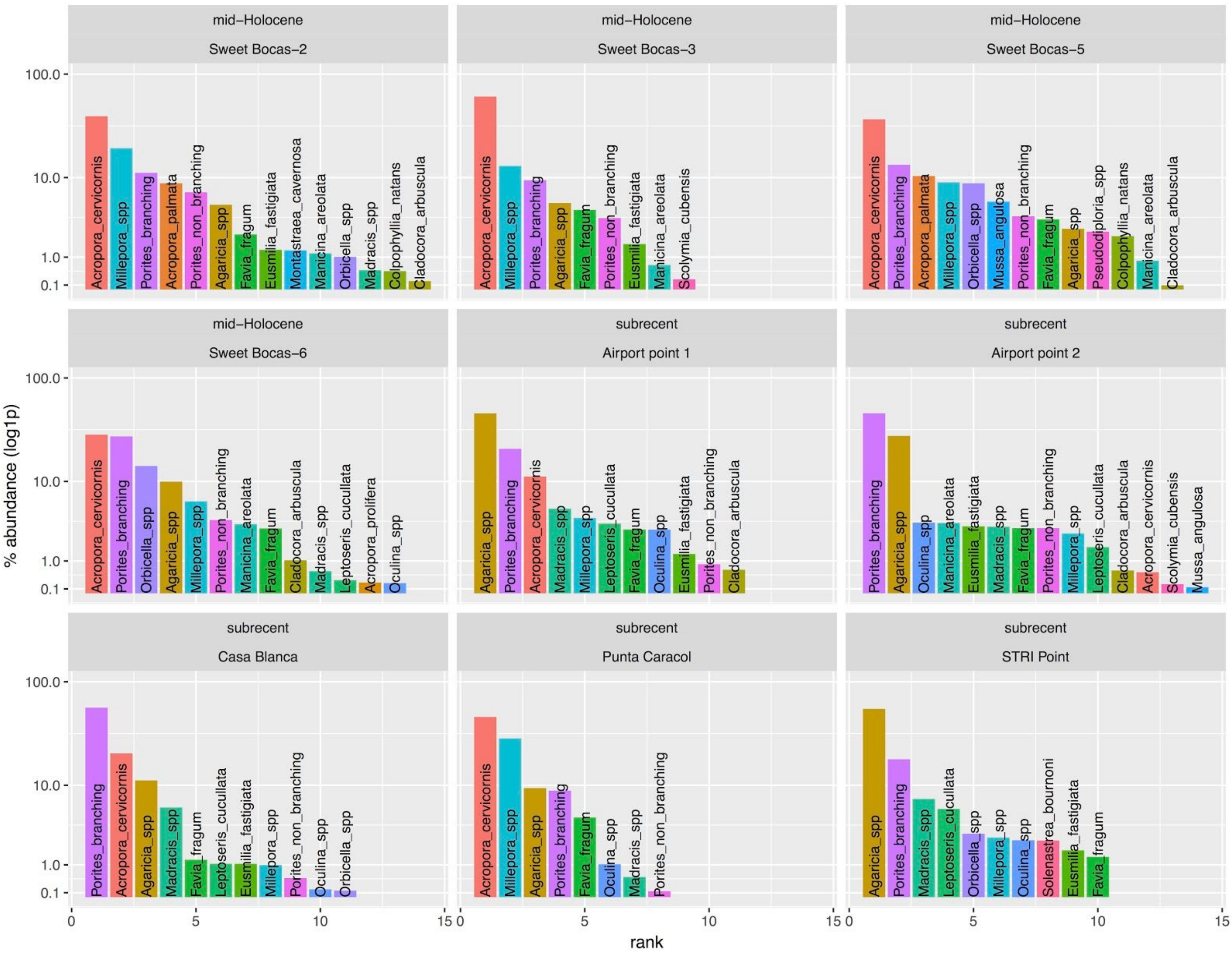


Figure S4. Rank-abundances (relative abundances based on mass of skeletal remains) of coral taxa in the modern reefs and mid-Holocene trenches.

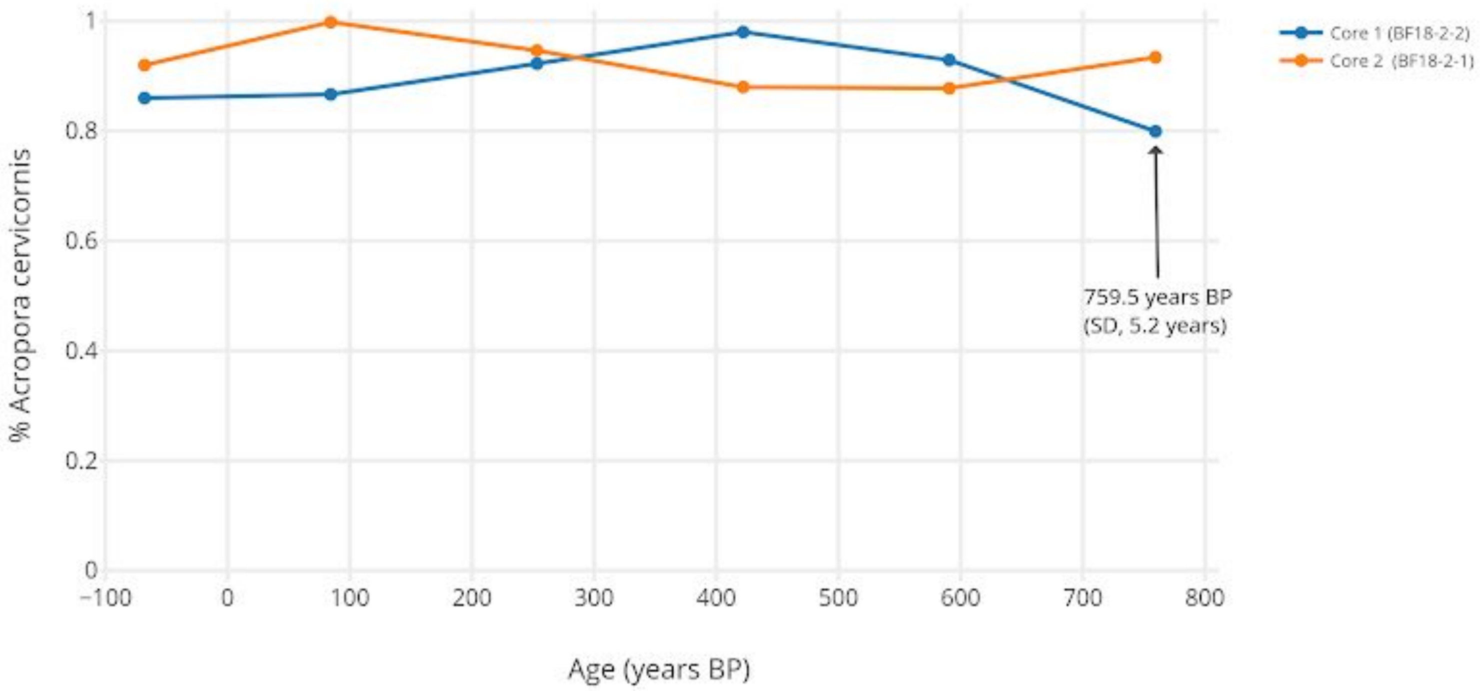


Figure S5. *Acropora cervicornis* dominated the reef at Punta Caracol over the last ~800 years. Relative abundance (proportion mass) of *A. cervicornis* to all other corals (including *Millepora*) in two cores taken ~2 m apart on the reef at Punta Caracol at 2.2 m water depth. Both cores included living coral at their tops. Chronology was estimated by dating a coral piece using U-Th (759.5 yBP) in the first core and assuming linear accretion rate to present day (2019) based on living coral at the top of the core.

**Table S1. Radiometric dates used in the study.** U-Th and calibrated-radiocarbon dating from coral pieces provide age control on modern and fossil samples to determine extent of time averaging (top), estimate reef accumulation rates at Punta Caracol reef (middle) and determine the age of the mid-Holocene reef (bottom). Almost all coral skeletons were chemically original with no calcite as observed from x-ray diffraction [11], and preservation generally exquisite with little bioerosion. * = mean taken from three measurements on same coral.

| Sample number | Site | Type | Method | medium | Age (years BP) | Year AD | 95% CI (years) | Ref. |
|---|---|---|---|---|---|---|---|---|
| To determine extent of time averaging in modern assemblages | | | | | | | | |
| AT13-10-1-1 | Punta Caracol | Bulk | U-Th | *Acropora cervicornis* | -58.1 | 2008 | 1.7 | This study |
| AT13-10-2-1 | Punta Caracol | Bulk | U-Th | *Acropora cervicornis* | -62.3 | 2012 | 1.2 | This study |
| AT13-10-2-1 | Punta Caracol | Bulk | U-Th | *Acropora cervicornis* | -58.2 | 2008 | 1.5 | This study |
| AT13-10-3-1 | Punta Caracol | Bulk | U-Th | *Acropora cervicornis* | -57.2 | 2007 | 1.4 | This study |
| AT13-10-4-1 | Punta Caracol | Bulk | U-Th | *Acropora cervicornis* | -52.5 | 2002 | 2.1 | This study |
| AT13-10-5-1 | Punta Caracol | Bulk | U-Th | *Acropora cervicornis* | -62.8 | 2012 | 1.3 | This study |
| VC13-4-2-[0-5] | Cayo Adriana | Core top | U-Th | *Acropora cervicornis* | 24 | 1926 | NA | [12] |
| VC13-3-1-[0-5] | Punto Donato | Core top | U-Th | *Acropora cervicornis* | -34 | 1984 | NA | [12] |
| VC13-3-1-[0-5] | Punto Donato | Core top | U-Th | *Acropora cervicornis* | -6 | 1956 | NA | [12] |
| To estimate reef chronology at Punta Caracol | | | | | | | | |
| BF18-2-2-[40-45] | Punta Caracol | Core | U-Th | *Acropora cervicornis* | 759.5 | 1190 | 5.2 | This study |
| To determine age of reefs in mid-Holocene site | | | | | | | | |
| AFH12-1a | Sunset Point | Hand | U-Th | Coral head | 7187 | | 8 | [11] |
| AFH12-1b | Sunset Point | Hand | U-Th | Coral head | 6708 | | 8 | [11] |
| AFH12-1c | Sunset Point | Hand | U-Th | Coral head | 6702* | | 42 | [11] |
| AFH12-1d | Sunset Point | Hand | U-Th | Coral head | 6671 | | 3 | [11] |
| AFH12-1e | Sunset Point | Hand | U-Th | Coral head | 6021 | | 3 | [11] |
| AFH12-1f | Sunset Point | Hand | U-Th | Coral head | 5834* | | 5 | [11] |
| AFH12-1g | Sunset Point | Hand | U-Th | Coral head | 5818 | | 7 | [11] |
| AFH12-1h | Sunset Point | Hand | U-Th | Coral head | 5781* | | 12 | [11] |
| AFH12-1i | Sunset Point | Hand | U-Th | Coral head | 5711 | | 3 | [11] |
| AT13-2-8 | Sweet Bocas | Bulk | U-Th | *Acropora cervicornis* | 6719 | | 13.3 | This study |
| AT13-3-11 | Sweet Bocas | Bulk | U-Th | *Acropora cervicornis* | 6982 | | 21.5 | This study |
| AT13-51A | Sweet Bocas | Hand | $^{14}C$ | *Acropora palmata* | 6638 | | - | This study |
| AT13-51A | Sweet Bocas | Hand | $^{14}C$ | *Acropora palmata* | 6638 | | - | This study |
| AT13-52A | Sweet Bocas | Hand | $^{14}C$ | *Acropora palmata* | 6589 | | - | This study |
| AT13-52A | Sweet Bocas | Hand | $^{14}C$ | *Acropora palmata* | 6589 | | - | This study |
| AT13-53A | Sweet Bocas | Hand | $^{14}C$ | *Acropora palmata* | 6626 | | - | This study |
| AT13-53A | Sweet Bocas | Hand | $^{14}C$ | *Acropora palmata* | 6626 | | - | This study |
| AT13-56A | Sweet Bocas | Hand | $^{14}C$ | *Acropora palmata* | 6533 | | - | This study |
| AT13-56A | Sweet Bocas | Hand | $^{14}C$ | *Acropora palmata* | 6533 | | - | This study |

**Table S2. Data and locality information of samples used in this study.** Raw data of coral taxa abundances (mass) in mid-Holocene and modern samples from Almirante Bay, Bocas del Toro.

| observ_order | sample | Age | Reef | Long | Lat | Depth (m) | genus | species | mass_g |
|---|---|---|---|---|---|---|---|---|---|
| MC_16 | AT12-4-3-2 | Sub Recent | SW Bocas Island | -82.25418 | 9.33184 | 6 | Porites | Porites spp_dfp | 0.51 |
| MC_16 | AT12-4-3-2 | Sub Recent | SW Bocas Island | -82.25418 | 9.33184 | 6 | Unknown | unknown | 41.51 |
| MC_16 | AT12-4-3-2 | Sub Recent | SW Bocas Island | -82.25418 | 9.33184 | 6 | Agaricia | Agaricia spp | 51.05 |
| MC_15 | AT12-4-5-1 | Sub Recent | SW Bocas Island | -82.25418 | 9.33184 | 10 | Acropora | Acropora cervicornis | 0.24 |
| MC_15 | AT12-4-5-1 | Sub Recent | SW Bocas Island | -82.25418 | 9.33184 | 10 | Madracis | Madracis spp | 1.06 |
| MC_15 | AT12-4-5-1 | Sub Recent | SW Bocas Island | -82.25418 | 9.33184 | 10 | Oculina | Oculina spp | 0.65 |
| MC_15 | AT12-4-5-1 | Sub Recent | SW Bocas Island | -82.25418 | 9.33184 | 10 | Porites | Porites spp_dfp | 0.84 |
| MC_15 | AT12-4-5-1 | Sub Recent | SW Bocas Island | -82.25418 | 9.33184 | 10 | Favia | Favia fragum | 1.59 |
| MC_15 | AT12-4-5-1 | Sub Recent | SW Bocas Island | -82.25418 | 9.33184 | 10 | Leptoseris | Leptoseris cucullata | 2.19 |
| MC_15 | AT12-4-5-1 | Sub Recent | SW Bocas Island | -82.25418 | 9.33184 | 10 | | other not coral | 4.64 |
| MC_15 | AT12-4-5-1 | Sub Recent | SW Bocas Island | -82.25418 | 9.33184 | 10 | Agaricia | Agaricia spp | 36.86 |
| MC_15 | AT12-4-5-1 | Sub Recent | SW Bocas Island | -82.25418 | 9.33184 | 10 | Unknown | unknown | 44.93 |
| ML_8 | AT13-10-1-1 | Sub Recent | Punta Caracol | -82.30383333 | 9.37845 | 1.9 | Oculina | Oculina diffusa | 0.14 |
| ML_8 | AT13-10-1-1 | Sub Recent | Punta Caracol | -82.30383333 | 9.37845 | 1.9 | Agaricia | Agaricia fragilis | 0.66 |
| ML_8 | AT13-10-1-1 | Sub Recent | Punta Caracol | -82.30383333 | 9.37845 | 1.9 | Favia | Favia fragum | 1.88 |
| ML_9 | AT13-10-2-1 | Sub Recent | Punta Caracol | -82.30383333 | 9.37845 | 1.9 | Porites | Porites colonensis | 0.10 |
| ML_8 | AT13-10-1-1 | Sub Recent | Punta Caracol | -82.30383333 | 9.37845 | 1.9 | Porites | Porites spp_dfp | 3.68 |
| ML_8 | AT13-10-1-1 | Sub Recent | Punta Caracol | -82.30383333 | 9.37845 | 1.9 | Agaricia | Agaricia agaricites | 9.58 |
| ML_8 | AT13-10-1-1 | Sub Recent | Punta Caracol | -82.30383333 | 9.37845 | 1.9 | Millepora | Millepora spp | 148.95 |
| ML_8 | AT13-10-1-1 | Sub Recent | Punta Caracol | -82.30383333 | 9.37845 | 1.9 | Acropora | Acropora cervicornis | 228.69 |
| ML_8 | AT13-10-1-1 | Sub Recent | Punta Caracol | -82.30383333 | 9.37845 | 1.9 | | other not coral | 494.30 |
| ML_9 | AT13-10-2-1 | Sub Recent | Punta Caracol | -82.30383333 | 9.37845 | 1.9 | Agaricia | Agaricia humilis | 9.31 |
| ML_9 | AT13-10-2-1 | Sub Recent | Punta Caracol | -82.30383333 | 9.37845 | 1.9 | Porites | Porites spp_dfp | 4.10 |
| ML_9 | AT13-10-2-1 | Sub Recent | Punta Caracol | -82.30383333 | 9.37845 | 1.9 | Acropora | Acropora cervicornis | 590.90 |
| ML_9 | AT13-10-2-1 | Sub Recent | Punta Caracol | -82.30383333 | 9.37845 | 1.9 | | other not coral | 412.10 |
| ML_9 | AT13-10-2-1 | Sub Recent | Punta Caracol | -82.30383333 | 9.37845 | 1.9 | Unknown | unknown | 140.09 |
| ML_9 | AT13-10-2-1 | Sub Recent | Punta Caracol | -82.30383333 | 9.37845 | 1.9 | Millepora | Millepora spp | 72.38 |
| ML_9 | AT13-10-2-1 | Sub Recent | Punta Caracol | -82.30383333 | 9.37845 | 1.9 | Agaricia | Agaricia lamarcki | 1.72 |
| ML_2 | AT13-2-10- | Holocene | Sweet Bocas Trench 1 | -82.27153333 | 9.360233333 | URS | Porites | Porites divaricata | 17.65 |
| ML_9 | AT13-10-2-1 | Sub Recent | Punta Caracol | -82.30383333 | 9.37845 | 1.9 | Agaricia | Agaricia agaricites | 0.63 |
| ML_10 | AT13-10-3-1 | Sub Recent | Punta Caracol | -82.30383333 | 9.37845 | 1.8 | Porites | Porites spp_dfp | 0.18 |
| ML_9 | AT13-10-2-1 | Sub Recent | Punta Caracol | -82.30383333 | 9.37845 | 1.9 | Madracis | Madracis spp | 0.19 |
| ML_10 | AT13-10-3-1 | Sub Recent | Punta Caracol | -82.30383333 | 9.37845 | 1.8 | Acropora | Acropora cervicornis | 259.74 |
| NM_5 | AT13-10-4-1 | Sub Recent | Punta Caracol | -82.30383333 | 9.37845 | 1.8 | Agaricia | Agaricia tenuifolia | 0.23 |
| NM_5 | AT13-10-4-1 | Sub Recent | Punta Caracol | -82.30383333 | 9.37845 | 1.8 | Favia | Favia fragum | 5.17 |
| ML_10 | AT13-10-3-1 | Sub Recent | Punta Caracol | -82.30383333 | 9.37845 | 1.8 | | other not coral | 1008.70 |
| NM_5 | AT13-10-4-1 | Sub Recent | Punta Caracol | -82.30383333 | 9.37845 | 1.8 | Agaricia | Agaricia lamarcki | 0.22 |
| NM_5 | AT13-10-4-1 | Sub Recent | Punta Caracol | -82.30383333 | 9.37845 | 1.8 | Agaricia | Agaricia agaricites | 3.21 |
| NM_5 | AT13-10-4-1 | Sub Recent | Punta Caracol | -82.30383333 | 9.37845 | 1.8 | Porites | Porites spp_dfp | 4.46 |
| NM_5 | AT13-10-4-1 | Sub Recent | Punta Caracol | -82.30383333 | 9.37845 | 1.8 | Agaricia | Agaricia fragilis | 1.39 |
| ML_10 | AT13-10-3-1 | Sub Recent | Punta Caracol | -82.30383333 | 9.37845 | 1.8 | Agaricia | Agaricia agaricites | 1.60 |
| ML_10 | AT13-10-3-1 | Sub Recent | Punta Caracol | -82.30383333 | 9.37845 | 1.8 | Madracis | Madracis decactis | 0.42 |
| ML_10 | AT13-10-3-1 | Sub Recent | Punta Caracol | -82.30383333 | 9.37845 | 1.8 | Agaricia | Agaricia spp | 1.75 |
| ML_10 | AT13-10-3-1 | Sub Recent | Punta Caracol | -82.30383333 | 9.37845 | 1.8 | Agaricia | Agaricia lamarcki | 3.42 |
| ML_2 | AT13-2-10- | Holocene | Sweet Bocas Trench 1 | -82.27153333 | 9.360233333 | URS | Unknown | unknown | 45.30 |
| ML_10 | AT13-10-3-1 | Sub Recent | Punta Caracol | -82.30383333 | 9.37845 | 1.8 | Millepora | Millepora spp | 224.45 |
| ML_10 | AT13-10-3-1 | Sub Recent | Punta Caracol | -82.30383333 | 9.37845 | 1.8 | Agaricia | Agaricia humilis | 2.84 |
| ML_10 | AT13-10-3-1 | Sub Recent | Punta Caracol | -82.30383333 | 9.37845 | 1.8 | Unknown | unknown | 218.53 |
| ML_10 | AT13-10-3-1 | Sub Recent | Punta Caracol | -82.30383333 | 9.37845 | 1.8 | Oculina | Oculina diffusa | 0.40 |
| ML_10 | AT13-10-3-1 | Sub Recent | Punta Caracol | -82.30383333 | 9.37845 | 1.8 | Favia | Favia fragum | 2.53 |
| ML_9 | AT13-10-2-1 | Sub Recent | Punta Caracol | -82.30383333 | 9.37845 | 1.9 | Favia | Favia fragum | 6.65 |
| ML_8 | AT13-10-1-1 | Sub Recent | Punta Caracol | -82.30383333 | 9.37845 | 1.9 | Unknown | unknown | 97.79 |

| | | | | | | | | | |
|---|---|---|---|---|---|---|---|---|---|
| NM_5 | AT13-10-4-1 | Sub Recent | Punta Caracol | -82.30383333 | 9.37845 | 1.8 | Agaricia | Agaricia humilis | 14.13 |
| NM_5 | AT13-10-4-1 | Sub Recent | Punta Caracol | -82.30383333 | 9.37845 | 1.8 | Unknown | unknown | 233.15 |
| NM_6 | AT13-10-5-1 | Sub Recent | Punta Caracol | -82.30383333 | 9.37845 | 1.7 | Agaricia | Agaricia agaricites | 4.95 |
| NM_6 | AT13-10-5-1 | Sub Recent | Punta Caracol | -82.30383333 | 9.37845 | 1.7 | Agaricia | Agaricia undata | 1.23 |
| NM_5 | AT13-10-4-1 | Sub Recent | Punta Caracol | -82.30383333 | 9.37845 | 1.8 | | other not coral | 309.86 |
| NM_6 | AT13-10-5-1 | Sub Recent | Punta Caracol | -82.30383333 | 9.37845 | 1.7 | Agaricia | Agaricia lamarcki | 1.60 |
| NM_6 | AT13-10-5-1 | Sub Recent | Punta Caracol | -82.30383333 | 9.37845 | 1.7 | Agaricia | Agaricia humilis | 3.17 |
| NM_5 | AT13-10-4-1 | Sub Recent | Punta Caracol | -82.30383333 | 9.37845 | 1.8 | Millepora | Millepora spp | 282.71 |
| NM_5 | AT13-10-4-1 | Sub Recent | Punta Caracol | -82.30383333 | 9.37845 | 1.8 | Agaricia | Agaricia spp | 42.00 |
| NM_6 | AT13-10-5-1 | Sub Recent | Punta Caracol | -82.30383333 | 9.37845 | 1.7 | Oculina | Oculina spp | 1.76 |
| NM_5 | AT13-10-4-1 | Sub Recent | Punta Caracol | -82.30383333 | 9.37845 | 1.8 | Acropora | Acropora cervicornis | 513.83 |
| NM_6 | AT13-10-5-1 | Sub Recent | Punta Caracol | -82.30383333 | 9.37845 | 1.7 | Porites | Porites spp_dfp | 44.44 |
| NM_6 | AT13-10-5-1 | Sub Recent | Punta Caracol | -82.30383333 | 9.37845 | 1.7 | Millepora | Millepora spp | 176.36 |
| NM_6 | AT13-10-5-1 | Sub Recent | Punta Caracol | -82.30383333 | 9.37845 | 1.7 | Unknown | unknown | 152.04 |
| NM_6 | AT13-10-5-1 | Sub Recent | Punta Caracol | -82.30383333 | 9.37845 | 1.7 | Acropora | Acropora cervicornis | 393.87 |
| ML_1 | AT13-2-1- | Holocene | Sweet Bocas Trench 1 | -82.27153333 | 9.360233333 | URS | Madracis | Madracis decactis | 4.49 |
| NM_6 | AT13-10-5-1 | Sub Recent | Punta Caracol | -82.30383333 | 9.37845 | 1.7 | Favia | Favia fragum | 5.68 |
| NM_6 | AT13-10-5-1 | Sub Recent | Punta Caracol | -82.30383333 | 9.37845 | 1.7 | | other not coral | 582.93 |
| ML_1 | AT13-2-1- | Holocene | Sweet Bocas Trench 1 | -82.27153333 | 9.360233333 | URS | Agaricia | Agaricia agaricites | 0.38 |
| ML_1 | AT13-2-1- | Holocene | Sweet Bocas Trench 1 | -82.27153333 | 9.360233333 | URS | Acropora | Acropora palmata | 1153.50 |
| ML_1 | AT13-2-1- | Holocene | Sweet Bocas Trench 1 | -82.27153333 | 9.360233333 | URS | Porites | Porites colonensis | 134.79 |
| ML_1 | AT13-2-1- | Holocene | Sweet Bocas Trench 1 | -82.27153333 | 9.360233333 | URS | Montastraea | Montastraea cavernosa | 29.49 |
| ML_1 | AT13-2-1- | Holocene | Sweet Bocas Trench 1 | -82.27153333 | 9.360233333 | URS | Manicina | Manicina areolata | 9.58 |
| ML_1 | AT13-2-1- | Holocene | Sweet Bocas Trench 1 | -82.27153333 | 9.360233333 | URS | cup | cup coral | 0.34 |
| ML_1 | AT13-2-1- | Holocene | Sweet Bocas Trench 1 | -82.27153333 | 9.360233333 | URS | Porites | Porites furcata | 24.27 |
| ML_1 | AT13-2-1- | Holocene | Sweet Bocas Trench 1 | -82.27153333 | 9.360233333 | URS | Porites | Porites divaricata | 3.90 |
| ML_1 | AT13-2-1- | Holocene | Sweet Bocas Trench 1 | -82.27153333 | 9.360233333 | URS | Favia | Favia fragum | 5.51 |
| ML_1 | AT13-2-1- | Holocene | Sweet Bocas Trench 1 | -82.27153333 | 9.360233333 | URS | Agaricia | Agaricia lamarcki | 13.59 |
| ML_1 | AT13-2-1- | Holocene | Sweet Bocas Trench 1 | -82.27153333 | 9.360233333 | URS | Eusmilia | Eusmilia fastigiata | 30.59 |
| ML_1 | AT13-2-1- | Holocene | Sweet Bocas Trench 1 | -82.27153333 | 9.360233333 | URS | | other not coral | 388.17 |
| ML_1 | AT13-2-1- | Holocene | Sweet Bocas Trench 1 | -82.27153333 | 9.360233333 | URS | Unknown | unknown | 527.80 |
| ML_1 | AT13-2-1- | Holocene | Sweet Bocas Trench 1 | -82.27153333 | 9.360233333 | URS | Acropora | Acropora cervicornis | 653.50 |
| ML_2 | AT13-2-10- | Holocene | Sweet Bocas Trench 1 | -82.27153333 | 9.360233333 | URS | Agaricia | Agaricia fragilis | 12.77 |
| ML_2 | AT13-2-10- | Holocene | Sweet Bocas Trench 1 | -82.27153333 | 9.360233333 | URS | Cladocora | Cladocora arbuscula | 0.21 |
| ML_2 | AT13-2-10- | Holocene | Sweet Bocas Trench 1 | -82.27153333 | 9.360233333 | URS | Porites | Porites spp_dfp | 11.01 |
| ML_2 | AT13-2-10- | Holocene | Sweet Bocas Trench 1 | -82.27153333 | 9.360233333 | URS | Agaricia | Agaricia agaricites | 11.19 |
| ML_2 | AT13-2-10- | Holocene | Sweet Bocas Trench 1 | -82.27153333 | 9.360233333 | URS | Millepora | Millepora spp | 488.11 |
| ML_2 | AT13-2-10- | Holocene | Sweet Bocas Trench 1 | -82.27153333 | 9.360233333 | URS | Acropora | Acropora cervicornis | 629.20 |
| ML_3 | AT13-2-8- | Holocene | Sweet Bocas Trench 1 | -82.27153333 | 9.360233333 | URS | Agaricia | Agaricia humilis | 0.30 |
| ML_3 | AT13-2-8- | Holocene | Sweet Bocas Trench 1 | -82.27153333 | 9.360233333 | URS | Agaricia | Agaricia lamarcki | 1.42 |
| ML_3 | AT13-2-8- | Holocene | Sweet Bocas Trench 1 | -82.27153333 | 9.360233333 | URS | Montastraea | Montastraea annularis | 9.09 |
| ML_3 | AT13-2-8- | Holocene | Sweet Bocas Trench 1 | -82.27153333 | 9.360233333 | URS | Porites | Porites divaricata | 1.55 |
| ML_3 | AT13-2-8- | Holocene | Sweet Bocas Trench 1 | -82.27153333 | 9.360233333 | URS | Porites | Porites furcata | 14.55 |
| ML_3 | AT13-2-8- | Holocene | Sweet Bocas Trench 1 | -82.27153333 | 9.360233333 | URS | Porites | Porites astreoides | 15.38 |
| ML_3 | AT13-2-8- | Holocene | Sweet Bocas Trench 1 | -82.27153333 | 9.360233333 | URS | Favia | Favia fragum | 16.63 |
| ML_3 | AT13-2-8- | Holocene | Sweet Bocas Trench 1 | -82.27153333 | 9.360233333 | URS | Acropora | Acropora cervicornis | 2006.60 |
| ML_3 | AT13-2-8- | Holocene | Sweet Bocas Trench 1 | -82.27153333 | 9.360233333 | URS | Manicina | Manicina areolata | 1.56 |
| ML_3 | AT13-2-8- | Holocene | Sweet Bocas Trench 1 | -82.27153333 | 9.360233333 | URS | | other not coral | 4.32 |
| ML_3 | AT13-2-8- | Holocene | Sweet Bocas Trench 1 | -82.27153333 | 9.360233333 | URS | Millepora | Millepora spp | 71.46 |
| ML_3 | AT13-2-8- | Holocene | Sweet Bocas Trench 1 | -82.27153333 | 9.360233333 | URS | Unknown | unknown | 313.62 |
| ML_4 | AT13-2-9- | Holocene | Sweet Bocas Trench 1 | -82.27153333 | 9.360233333 | URS | Favia | Favia fragum | 0.87 |
| ML_4 | AT13-2-9- | Holocene | Sweet Bocas Trench 1 | -82.27153333 | 9.360233333 | URS | Agaricia | Agaricia lamarcki | 11.44 |
| ML_4 | AT13-2-9- | Holocene | Sweet Bocas Trench 1 | -82.27153333 | 9.360233333 | URS | Porites | Porites divaricata | 15.50 |
| ML_4 | AT13-2-9- | Holocene | Sweet Bocas Trench 1 | -82.27153333 | 9.360233333 | URS | Agaricia | Agaricia agaricites | 17.26 |
| ML_4 | AT13-2-9- | Holocene | Sweet Bocas Trench 1 | -82.27153333 | 9.360233333 | URS | Porites | Porites colonensis | 118.17 |
| ML_4 | AT13-2-9- | Holocene | Sweet Bocas Trench 1 | -82.27153333 | 9.360233333 | URS | Porites | Porites spp_dfp | 147.40 |
| ML_4 | AT13-2-9- | Holocene | Sweet Bocas Trench 1 | -82.27153333 | 9.360233333 | URS | Unknown | unknown | 416.54 |

| | | | | | | | | | |
|---|---|---|---|---|---|---|---|---|---|
| ML_4 | AT13-2-9- | Holocene | Sweet Bocas Trench 1 | -82.27153333 | 9.360233333 | URS | Millepora | Millepora spp | 620.30 |
| ML_4 | AT13-2-9- | Holocene | Sweet Bocas Trench 1 | -82.27153333 | 9.360233333 | URS | Acropora | Acropora cervicornis | 638.85 |
| ML_5 | AT13-3-10- | Holocene | Sweet Bocas Trench 2 | -82.27143333 | 9.359933333 | URS | Agaricia | Agaricia tenuifolia | 0.28 |
| ML_5 | AT13-3-10- | Holocene | Sweet Bocas Trench 2 | -82.27143333 | 9.359933333 | URS | Agaricia | Agaricia humilis | 14.07 |
| ML_5 | AT13-3-10- | Holocene | Sweet Bocas Trench 2 | -82.27143333 | 9.359933333 | URS | Porites | Porites colonensis | 12.84 |
| ML_5 | AT13-3-10- | Holocene | Sweet Bocas Trench 2 | -82.27143333 | 9.359933333 | URS | Porites | Porites divaricata | 20.82 |
| ML_5 | AT13-3-10- | Holocene | Sweet Bocas Trench 2 | -82.27143333 | 9.359933333 | URS | | other not coral | 21.90 |
| ML_5 | AT13-3-10- | Holocene | Sweet Bocas Trench 2 | -82.27143333 | 9.359933333 | URS | Favia | Favia fragum | 25.54 |
| ML_5 | AT13-3-10- | Holocene | Sweet Bocas Trench 2 | -82.27143333 | 9.359933333 | URS | Millepora | Millepora spp | 76.72 |
| MC_4 | AT12-1-1-1 | Sub Recent | STRI Point | -82.262555 | 9.349446 | 12 | Madracis | Madracis spp | 0.12 |
| MC_4 | AT12-1-1-1 | Sub Recent | STRI Point | -82.262555 | 9.349446 | 12 | Oculina | Oculina diffusa | 0.14 |
| MC_4 | AT12-1-1-1 | Sub Recent | STRI Point | -82.262555 | 9.349446 | 12 | Favia | Favia fragum | 0.24 |
| MC_4 | AT12-1-1-1 | Sub Recent | STRI Point | -82.262555 | 9.349446 | 12 | Eusmilia | Eusmilia fastigiata | 0.38 |
| MC_4 | AT12-1-1-1 | Sub Recent | STRI Point | -82.262555 | 9.349446 | 12 | Solenastrea | Solenastrea bournoni | 0.71 |
| MC_4 | AT12-1-1-1 | Sub Recent | STRI Point | -82.262555 | 9.349446 | 12 | Leptoseris | Leptoseris cucullata | 0.72 |
| MC_4 | AT12-1-1-1 | Sub Recent | STRI Point | -82.262555 | 9.349446 | 12 | Millepora | Millepora spp | 0.85 |
| MC_4 | AT12-1-1-1 | Sub Recent | STRI Point | -82.262555 | 9.349446 | 12 | Porites | Porites spp_dfp | 2.72 |
| MC_4 | AT12-1-1-1 | Sub Recent | STRI Point | -82.262555 | 9.349446 | 12 | | other not coral | 27.28 |
| MC_4 | AT12-1-1-1 | Sub Recent | STRI Point | -82.262555 | 9.349446 | 12 | Agaricia | Agaricia spp | 33.62 |
| MC_4 | AT12-1-1-1 | Sub Recent | STRI Point | -82.262555 | 9.349446 | 12 | Unknown | unknown | 69.19 |
| NM_15 | AT12-1-2-1 | Sub Recent | STRI Point | -82.262555 | 9.349446 | 10 | Madracis | Madracis asperula | 1.89 |
| NM_15 | AT12-1-2-1 | Sub Recent | STRI Point | -82.262555 | 9.349446 | 10 | Leptoseris | Leptoseris cucullata | 0.34 |
| NM_15 | AT12-1-2-1 | Sub Recent | STRI Point | -82.262555 | 9.349446 | 10 | Orbicella | Orbicella spp | 0.36 |
| NM_15 | AT12-1-2-1 | Sub Recent | STRI Point | -82.262555 | 9.349446 | 10 | Oculina | Oculina spp | 0.45 |
| NM_15 | AT12-1-2-1 | Sub Recent | STRI Point | -82.262555 | 9.349446 | 10 | Madracis | Madracis decactis | 1.35 |
| NM_15 | AT12-1-2-1 | Sub Recent | STRI Point | -82.262555 | 9.349446 | 10 | | other not coral | 2.14 |
| NM_15 | AT12-1-2-1 | Sub Recent | STRI Point | -82.262555 | 9.349446 | 10 | Porites | Porites spp_dfp | 6.03 |
| NM_15 | AT12-1-2-1 | Sub Recent | STRI Point | -82.262555 | 9.349446 | 10 | Unknown | unknown | 46.89 |
| NM_15 | AT12-1-2-1 | Sub Recent | STRI Point | -82.262555 | 9.349446 | 10 | Agaricia | Agaricia spp | 112.15 |
| MC_13 | AT12-1-3-2 | Sub Recent | STRI Point | -82.262555 | 9.349446 | 8 | | other not coral | 0.67 |
| MC_13 | AT12-1-3-2 | Sub Recent | STRI Point | -82.262555 | 9.349446 | 8 | Orbicella | Orbicella spp | 0.94 |
| MC_13 | AT12-1-3-2 | Sub Recent | STRI Point | -82.262555 | 9.349446 | 8 | Leptoseris | Leptoseris cucullata | 1.42 |
| MC_13 | AT12-1-3-2 | Sub Recent | STRI Point | -82.262555 | 9.349446 | 8 | Madracis | Madracis decactis | 2.29 |
| MC_13 | AT12-1-3-2 | Sub Recent | STRI Point | -82.262555 | 9.349446 | 8 | unknown | unknown | 10.16 |
| MC_13 | AT12-1-3-2 | Sub Recent | STRI Point | -82.262555 | 9.349446 | 8 | Porites | Porites spp_dfp | 24.71 |
| MC_13 | AT12-1-3-2 | Sub Recent | STRI Point | -82.262555 | 9.349446 | 8 | Agaricia | Agaricia spp | 108.13 |
| MC_12 | AT12-4-1-2 | Sub Recent | SW Bocas Island | -82.25418 | 9.33184 | 2 | cup | cup coral | 0.13 |
| MC_12 | AT12-4-1-2 | Sub Recent | SW Bocas Island | -82.25418 | 9.33184 | 2 | Cladocora | Cladocora arbuscula | 0.20 |
| MC_12 | AT12-4-1-2 | Sub Recent | SW Bocas Island | -82.25418 | 9.33184 | 2 | | other not coral | 2.51 |
| MC_12 | AT12-4-1-2 | Sub Recent | SW Bocas Island | -82.25418 | 9.33184 | 2 | Millepora | Millepora spp | 6.86 |
| MC_12 | AT12-4-1-2 | Sub Recent | SW Bocas Island | -82.25418 | 9.33184 | 2 | Acropora | Acropora cervicornis | 11.86 |
| MC_12 | AT12-4-1-2 | Sub Recent | SW Bocas Island | -82.25418 | 9.33184 | 2 | Agaricia | Agaricia spp | 40.51 |
| MC_17 | AT12-4-2-1 | Sub Recent | SW Bocas Island | -82.25418 | 9.33184 | 4 | Oculina | Oculina spp | 3.93 |
| MC_17 | AT12-4-2-1 | Sub Recent | SW Bocas Island | -82.25418 | 9.33184 | 4 | Eusmilia | Eusmilia fastigiata | 4.51 |
| MC_17 | AT12-4-2-1 | Sub Recent | SW Bocas Island | -82.25418 | 9.33184 | 4 | Millepora | Millepora spp | 4.53 |
| MC_17 | AT12-4-2-1 | Sub Recent | SW Bocas Island | -82.25418 | 9.33184 | 4 | Madracis | Madracis spp | 13.86 |
| MC_17 | AT12-4-2-1 | Sub Recent | SW Bocas Island | -82.25418 | 9.33184 | 4 | Porites | Porites spp_dfp | 76.82 |
| MC_17 | AT12-4-2-1 | Sub Recent | SW Bocas Island | -82.25418 | 9.33184 | 4 | Acropora | Acropora cervicornis | 80.81 |
| MC_17 | AT12-4-2-1 | Sub Recent | SW Bocas Island | -82.25418 | 9.33184 | 4 | Agaricia | Agaricia spp | 89.00 |
| MC_17 | AT12-4-2-1 | Sub Recent | SW Bocas Island | -82.25418 | 9.33184 | 4 | Unknown | unknown | 279.71 |
| MC_16 | AT12-4-3-2 | Sub Recent | SW Bocas Island | -82.25418 | 9.33184 | 6 | Madracis | Madracis spp | 0.05 |
| MC_16 | AT12-4-3-2 | Sub Recent | SW Bocas Island | -82.25418 | 9.33184 | 6 | Acropora | Acropora cervicornis | 0.09 |
| ML_5 | AT13-3-10- | Holocene | Sweet Bocas Trench 2 | -82.27143333 | 9.359933333 | URS | Unknown | unknown | 410.97 |
| ML_5 | AT13-3-10- | Holocene | Sweet Bocas Trench 2 | -82.27143333 | 9.359933333 | URS | Acropora | Acropora cervicornis | 1395.57 |
| ML_6 | AT13-3-11- | Holocene | Sweet Bocas Trench 2 | -82.27143333 | 9.359933333 | URS | Agaricia | Agaricia grahamae | 24.29 |
| ML_6 | AT13-3-11- | Holocene | Sweet Bocas Trench 2 | -82.27143333 | 9.359933333 | URS | Porites | Porites astreoides | 24.75 |
| ML_6 | AT13-3-11- | Holocene | Sweet Bocas Trench 2 | -82.27143333 | 9.359933333 | URS | Acropora | Acropora cervicornis | 2128.77 |

| | | | | | | | | | |
|---|---|---|---|---|---|---|---|---|---|
| ML_6 | AT13-3-11- | Holocene | Sweet Bocas Trench 2 | -82.27143333 | 9.359933333 | URS | Scolymia | Scolymia cubensis | 1.03 |
| MC_12 | AT12-4-1-2 | Sub Recent | SW Bocas Island | -82.25418 | 9.33184 | 2 | Porites | Porites spp_dfp | 91.66 |
| MC_12 | AT12-4-1-2 | Sub Recent | SW Bocas Island | -82.25418 | 9.33184 | 2 | Unknown | unknown | 96.41 |
| MC_17 | AT12-4-2-1 | Sub Recent | SW Bocas Island | -82.25418 | 9.33184 | 4 | Cladocora | Cladocora arbuscula | 0.10 |
| MC_17 | AT12-4-2-1 | Sub Recent | SW Bocas Island | -82.25418 | 9.33184 | 4 | | other not coral | 0.16 |
| MC_17 | AT12-4-2-1 | Sub Recent | SW Bocas Island | -82.25418 | 9.33184 | 4 | Favia | Favia fragum | 0.25 |
| MC_17 | AT12-4-2-1 | Sub Recent | SW Bocas Island | -82.25418 | 9.33184 | 4 | Leptoseris | Leptoseris cucullata | 0.37 |
| MC_17 | AT12-4-2-1 | Sub Recent | SW Bocas Island | -82.25418 | 9.33184 | 4 | Porites | Porites astreoides | 1.92 |
| ML_6 | AT13-3-11- | Holocene | Sweet Bocas Trench 2 | -82.27143333 | 9.359933333 | URS | Porites | Porites divaricata | 3.57 |
| ML_6 | AT13-3-11- | Holocene | Sweet Bocas Trench 2 | -82.27143333 | 9.359933333 | URS | | other not coral | 7.40 |
| ML_6 | AT13-3-11- | Holocene | Sweet Bocas Trench 2 | -82.27143333 | 9.359933333 | URS | Favia | Favia fragum | 9.09 |
| ML_6 | AT13-3-11- | Holocene | Sweet Bocas Trench 2 | -82.27143333 | 9.359933333 | URS | Porites | Porites spp_dfp | 11.74 |
| ML_6 | AT13-3-11- | Holocene | Sweet Bocas Trench 2 | -82.27143333 | 9.359933333 | URS | Agaricia | Agaricia humilis | 13.24 |
| ML_6 | AT13-3-11- | Holocene | Sweet Bocas Trench 2 | -82.27143333 | 9.359933333 | URS | Eusmilia | Eusmilia fastigiata | 47.18 |
| ML_6 | AT13-3-11- | Holocene | Sweet Bocas Trench 2 | -82.27143333 | 9.359933333 | URS | Millepora | Millepora spp | 83.81 |
| ML_6 | AT13-3-11- | Holocene | Sweet Bocas Trench 2 | -82.27143333 | 9.359933333 | URS | Unknown | unknown | 258.49 |
| ML_7 | AT13-3-12- | Holocene | Sweet Bocas Trench 2 | -82.27143333 | 9.359933333 | URS | Porites | Porites astreoides | 0.40 |
| ML_7 | AT13-3-12- | Holocene | Sweet Bocas Trench 2 | -82.27143333 | 9.359933333 | URS | Agaricia | Agaricia lamarcki | 2.06 |
| ML_7 | AT13-3-12- | Holocene | Sweet Bocas Trench 2 | -82.27143333 | 9.359933333 | URS | | other not coral | 2.07 |
| ML_7 | AT13-3-12- | Holocene | Sweet Bocas Trench 2 | -82.27143333 | 9.359933333 | URS | Favia | Favia fragum | 6.21 |
| ML_7 | AT13-3-12- | Holocene | Sweet Bocas Trench 2 | -82.27143333 | 9.359933333 | URS | Agaricia | Agaricia humilis | 8.63 |
| ML_7 | AT13-3-12- | Holocene | Sweet Bocas Trench 2 | -82.27143333 | 9.359933333 | URS | Millepora | Millepora spp | 59.24 |
| ML_7 | AT13-3-12- | Holocene | Sweet Bocas Trench 2 | -82.27143333 | 9.359933333 | URS | Unknown | unknown | 282.66 |
| ML_7 | AT13-3-12- | Holocene | Sweet Bocas Trench 2 | -82.27143333 | 9.359933333 | URS | Acropora | Acropora cervicornis | 2093.71 |
| NM_1 | AT13-3-13- | Holocene | Sweet Bocas Trench 2 | -82.27143333 | 9.359933333 | URS | Agaricia | Agaricia lamarcki | 0.59 |
| NM_1 | AT13-3-13- | Holocene | Sweet Bocas Trench 2 | -82.27143333 | 9.359933333 | URS | Porites | Porites colonensis | 0.74 |
| NM_1 | AT13-3-13- | Holocene | Sweet Bocas Trench 2 | -82.27143333 | 9.359933333 | URS | Porites | Porites divaricata | 1.06 |
| NM_1 | AT13-3-13- | Holocene | Sweet Bocas Trench 2 | -82.27143333 | 9.359933333 | URS | Porites | Porites astreoides | 1.12 |
| NM_1 | AT13-3-13- | Holocene | Sweet Bocas Trench 2 | -82.27143333 | 9.359933333 | URS | Agaricia | Agaricia agaricites | 1.97 |
| NM_1 | AT13-3-13- | Holocene | Sweet Bocas Trench 2 | -82.27143333 | 9.359933333 | URS | Manicina | Manicina areolata | 5.52 |
| NM_1 | AT13-3-13- | Holocene | Sweet Bocas Trench 2 | -82.27143333 | 9.359933333 | URS | Favia | Favia fragum | 8.58 |
| NM_1 | AT13-3-13- | Holocene | Sweet Bocas Trench 2 | -82.27143333 | 9.359933333 | URS | | other not coral | 20.84 |
| NM_1 | AT13-3-13- | Holocene | Sweet Bocas Trench 2 | -82.27143333 | 9.359933333 | URS | Millepora | Millepora spp | 27.54 |
| NM_1 | AT13-3-13- | Holocene | Sweet Bocas Trench 2 | -82.27143333 | 9.359933333 | URS | Unknown | unknown | 376.92 |
| NM_1 | AT13-3-13- | Holocene | Sweet Bocas Trench 2 | -82.27143333 | 9.359933333 | URS | Acropora | Acropora cervicornis | 2722.64 |
| NM_2 | AT13-3-14- | Holocene | Sweet Bocas Trench 2 | -82.27143333 | 9.359933333 | URS | Agaricia | Agaricia tenuifolia | 0.12 |
| NM_2 | AT13-3-14- | Holocene | Sweet Bocas Trench 2 | -82.27143333 | 9.359933333 | URS | Agaricia | Agaricia spp | 1.31 |
| NM_2 | AT13-3-14- | Holocene | Sweet Bocas Trench 2 | -82.27143333 | 9.359933333 | URS | Porites | Porites colonensis | 1.43 |
| NM_2 | AT13-3-14- | Holocene | Sweet Bocas Trench 2 | -82.27143333 | 9.359933333 | URS | Favia | Favia fragum | 1.89 |
| NM_2 | AT13-3-14- | Holocene | Sweet Bocas Trench 2 | -82.27143333 | 9.359933333 | URS | | other not coral | 2.40 |
| NM_2 | AT13-3-14- | Holocene | Sweet Bocas Trench 2 | -82.27143333 | 9.359933333 | URS | Porites | Porites astreoides | 8.03 |
| NM_2 | AT13-3-14- | Holocene | Sweet Bocas Trench 2 | -82.27143333 | 9.359933333 | URS | Agaricia | Agaricia humilis | 9.81 |
| NM_2 | AT13-3-14- | Holocene | Sweet Bocas Trench 2 | -82.27143333 | 9.359933333 | URS | Agaricia | Agaricia agaricites | 10.50 |
| NM_2 | AT13-3-14- | Holocene | Sweet Bocas Trench 2 | -82.27143333 | 9.359933333 | URS | Porites | Porites furcata | 36.33 |
| NM_2 | AT13-3-14- | Holocene | Sweet Bocas Trench 2 | -82.27143333 | 9.359933333 | URS | Millepora | Millepora spp | 325.05 |
| NM_2 | AT13-3-14- | Holocene | Sweet Bocas Trench 2 | -82.27143333 | 9.359933333 | URS | Unknown | unknown | 507.26 |
| NM_2 | AT13-3-14- | Holocene | Sweet Bocas Trench 2 | -82.27143333 | 9.359933333 | URS | Porites | Porites spp_dfp | 616.30 |
| NM_2 | AT13-3-14- | Holocene | Sweet Bocas Trench 2 | -82.27143333 | 9.359933333 | URS | Acropora | Acropora cervicornis | 807.18 |
| NM_3 | AT13-5-1- | Holocene | Sweet Bocas Trench 4 | -82.2718 | 9.359983333 | URS | Colpophyllia | Colpophyllia natans | 27.25 |
| NM_3 | AT13-5-1- | Holocene | Sweet Bocas Trench 4 | -82.2718 | 9.359983333 | URS | Orbicella | Orbicella spp | 30.95 |
| NM_3 | AT13-5-1- | Holocene | Sweet Bocas Trench 4 | -82.2718 | 9.359983333 | URS | Mussa | Mussa angulosa | 281.06 |
| NM_3 | AT13-5-1- | Holocene | Sweet Bocas Trench 4 | -82.2718 | 9.359983333 | URS | Agaricia | Agaricia agaricites | 0.63 |
| NM_3 | AT13-5-1- | Holocene | Sweet Bocas Trench 4 | -82.2718 | 9.359983333 | URS | Porites | Porites furcata | 0.99 |
| NM_3 | AT13-5-1- | Holocene | Sweet Bocas Trench 4 | -82.2718 | 9.359983333 | URS | Agaricia | Agaricia grahamae | 1.12 |
| NM_3 | AT13-5-1- | Holocene | Sweet Bocas Trench 4 | -82.2718 | 9.359983333 | URS | Agaricia | Agaricia lamarcki | 1.15 |
| NM_3 | AT13-5-1- | Holocene | Sweet Bocas Trench 4 | -82.2718 | 9.359983333 | URS | Agaricia | Agaricia humilis | 3.16 |
| NM_3 | AT13-5-1- | Holocene | Sweet Bocas Trench 4 | -82.2718 | 9.359983333 | URS | | other not coral | 3.81 |

| | | | | | | | | | |
|---|---|---|---|---|---|---|---|---|---|
| NM_3 | AT13-5-1- | Holocene | Sweet Bocas Trench 4 | -82.2718 | 9.359983333 | URS | Manicina | Manicina areolata | 6.59 |
| NM_3 | AT13-5-1- | Holocene | Sweet Bocas Trench 4 | -82.2718 | 9.359983333 | URS | Porites | Porites spp_dfp | 8.34 |
| NM_3 | AT13-5-1- | Holocene | Sweet Bocas Trench 4 | -82.2718 | 9.359983333 | URS | Favia | Favia fragum | 17.89 |
| NM_3 | AT13-5-1- | Holocene | Sweet Bocas Trench 4 | -82.2718 | 9.359983333 | URS | Montastraea | Montastraea annularis | 24.36 |
| NM_3 | AT13-5-1- | Holocene | Sweet Bocas Trench 4 | -82.2718 | 9.359983333 | URS | Millepora | Millepora spp | 57.03 |
| NM_3 | AT13-5-1- | Holocene | Sweet Bocas Trench 4 | -82.2718 | 9.359983333 | URS | Montastraea | Montastraea franksi | 133.17 |
| NM_3 | AT13-5-1- | Holocene | Sweet Bocas Trench 4 | -82.2718 | 9.359983333 | URS | Unknown | unknown | 715.81 |
| NM_3 | AT13-5-1- | Holocene | Sweet Bocas Trench 4 | -82.2718 | 9.359983333 | URS | Acropora | Acropora cervicornis | 1477.43 |
| NM_4 | AT13-5-2- | Holocene | Sweet Bocas Trench 4 | -82.2718 | 9.359983333 | URS | Agaricia | Agaricia fragilis | 0.40 |
| NM_4 | AT13-5-2- | Holocene | Sweet Bocas Trench 4 | -82.2718 | 9.359983333 | URS | Agaricia | Agaricia agaricites | 0.59 |
| NM_4 | AT13-5-2- | Holocene | Sweet Bocas Trench 4 | -82.2718 | 9.359983333 | URS | Agaricia | Agaricia undata | 0.77 |
| NM_4 | AT13-5-2- | Holocene | Sweet Bocas Trench 4 | -82.2718 | 9.359983333 | URS | | other not coral | 1.79 |
| NM_4 | AT13-5-2- | Holocene | Sweet Bocas Trench 4 | -82.2718 | 9.359983333 | URS | Agaricia | Agaricia lamarcki | 4.92 |
| NM_4 | AT13-5-2- | Holocene | Sweet Bocas Trench 4 | -82.2718 | 9.359983333 | URS | Millepora | Millepora spp | 5.03 |
| NM_4 | AT13-5-2- | Holocene | Sweet Bocas Trench 4 | -82.2718 | 9.359983333 | URS | Favia | Favia fragum | 7.78 |
| NM_4 | AT13-5-2- | Holocene | Sweet Bocas Trench 4 | -82.2718 | 9.359983333 | URS | Porites | Porites spp_dfp | 289.06 |
| NM_4 | AT13-5-2- | Holocene | Sweet Bocas Trench 4 | -82.2718 | 9.359983333 | URS | Unknown | unknown | 410.37 |
| NM_4 | AT13-5-2- | Holocene | Sweet Bocas Trench 4 | -82.2718 | 9.359983333 | URS | Acropora | Acropora palmata | 615.90 |
| NM_4 | AT13-5-2- | Holocene | Sweet Bocas Trench 4 | -82.2718 | 9.359983333 | URS | Acropora | Acropora cervicornis | 812.87 |
| NM_12 | AT13-5-3- | Holocene | Sweet Bocas Trench 4 | -82.2718 | 9.359983333 | URS | Montastraea | Montastraea annularis | 329.87 |
| NM_12 | AT13-5-3- | Holocene | Sweet Bocas Trench 4 | -82.2718 | 9.359983333 | URS | Cladocora | Cladocora arbuscula | 0.17 |
| NM_12 | AT13-5-3- | Holocene | Sweet Bocas Trench 4 | -82.2718 | 9.359983333 | URS | | other not coral | 1.32 |
| NM_12 | AT13-5-3- | Holocene | Sweet Bocas Trench 4 | -82.2718 | 9.359983333 | URS | Agaricia | Agaricia spp | 11.42 |
| NM_12 | AT13-5-3- | Holocene | Sweet Bocas Trench 4 | -82.2718 | 9.359983333 | URS | Favia | Favia fragum | 14.57 |
| NM_12 | AT13-5-3- | Holocene | Sweet Bocas Trench 4 | -82.2718 | 9.359983333 | URS | Acropora | Acropora palmata | 14.64 |
| NM_12 | AT13-5-3- | Holocene | Sweet Bocas Trench 4 | -82.2718 | 9.359983333 | URS | Diploria | Diploria spp | 113.10 |
| NM_12 | AT13-5-3- | Holocene | Sweet Bocas Trench 4 | -82.2718 | 9.359983333 | URS | Porites | Porites colonensis | 10.69 |
| NM_12 | AT13-5-3- | Holocene | Sweet Bocas Trench 4 | -82.2718 | 9.359983333 | URS | Unknown | unknown | 367.28 |
| NM_12 | AT13-5-3- | Holocene | Sweet Bocas Trench 4 | -82.2718 | 9.359983333 | URS | Porites | Porites spp_dfp | 528.30 |
| NM_12 | AT13-5-3- | Holocene | Sweet Bocas Trench 4 | -82.2718 | 9.359983333 | URS | Millepora | Millepora spp | 569.38 |
| NM_12 | AT13-5-3- | Holocene | Sweet Bocas Trench 4 | -82.2718 | 9.359983333 | URS | Acropora | Acropora cervicornis | 1445.17 |
| NM_12 | AT13-5-3- | Holocene | Sweet Bocas Trench 4 | -82.2718 | 9.359983333 | URS | Porites | Porites astreoides | 261.21 |
| MC_8 | AT13-6-1- | Holocene | Sweet Bocas Trench 5 | -82.272318 | 9.359966667 | URS | Madracis | Madracis spp | 2.54 |
| MC_8 | AT13-6-1- | Holocene | Sweet Bocas Trench 5 | -82.272318 | 9.359966667 | URS | Favia | Favia fragum | 3.15 |
| MC_8 | AT13-6-1- | Holocene | Sweet Bocas Trench 5 | -82.272318 | 9.359966667 | URS | Montastraea | Montastraea annularis | 8.49 |
| MC_8 | AT13-6-1- | Holocene | Sweet Bocas Trench 5 | -82.272318 | 9.359966667 | URS | Manicina | Manicina areolata | 10.36 |
| MC_8 | AT13-6-1- | Holocene | Sweet Bocas Trench 5 | -82.272318 | 9.359966667 | URS | Porites | Porites astreoides | 39.84 |
| MC_8 | AT13-6-1- | Holocene | Sweet Bocas Trench 5 | -82.272318 | 9.359966667 | URS | Porites | Porites spp_dfp | 54.26 |
| MC_8 | AT13-6-1- | Holocene | Sweet Bocas Trench 5 | -82.272318 | 9.359966667 | URS | Millepora | Millepora spp | 57.12 |
| MC_8 | AT13-6-1- | Holocene | Sweet Bocas Trench 5 | -82.272318 | 9.359966667 | URS | Agaricia | Agaricia spp | 133.85 |
| MC_8 | AT13-6-1- | Holocene | Sweet Bocas Trench 5 | -82.272318 | 9.359966667 | URS | Unknown | unknown | 424.42 |
| MC_8 | AT13-6-1- | Holocene | Sweet Bocas Trench 5 | -82.272318 | 9.359966667 | URS | Acropora | Acropora cervicornis | 1906.23 |
| MC_9 | AT13-6-2- | Holocene | Sweet Bocas Trench 5 | -82.272318 | 9.359966667 | URS | Leptoseris | Leptoseris cucullata | 2.21 |
| MC_9 | AT13-6-2- | Holocene | Sweet Bocas Trench 5 | -82.272318 | 9.359966667 | URS | Cladocora | Cladocora arbuscula | 0.08 |
| MC_9 | AT13-6-2- | Holocene | Sweet Bocas Trench 5 | -82.272318 | 9.359966667 | URS | Porites | Porites astreoides | 0.22 |
| MC_9 | AT13-6-2- | Holocene | Sweet Bocas Trench 5 | -82.272318 | 9.359966667 | URS | Oculina | Oculina diffusa | 0.52 |
| MC_9 | AT13-6-2- | Holocene | Sweet Bocas Trench 5 | -82.272318 | 9.359966667 | URS | Acropora | Acropora prolifera | 1.411 |
| MC_9 | AT13-6-2- | Holocene | Sweet Bocas Trench 5 | -82.272318 | 9.359966667 | URS | Porites | Porites colonensis | 1.68 |
| MC_9 | AT13-6-2- | Holocene | Sweet Bocas Trench 5 | -82.272318 | 9.359966667 | URS | Manicina | Manicina areolata | 17.71 |
| MC_9 | AT13-6-2- | Holocene | Sweet Bocas Trench 5 | -82.272318 | 9.359966667 | URS | Favia | Favia fragum | 33.86 |
| MC_9 | AT13-6-2- | Holocene | Sweet Bocas Trench 5 | -82.272318 | 9.359966667 | URS | Agaricia | Agaricia spp | 74.62 |
| MC_9 | AT13-6-2- | Holocene | Sweet Bocas Trench 5 | -82.272318 | 9.359966667 | URS | Millepora | Millepora spp | 110.61 |
| MC_9 | AT13-6-2- | Holocene | Sweet Bocas Trench 5 | -82.272318 | 9.359966667 | URS | | other not coral | 264.19 |
| MC_9 | AT13-6-2- | Holocene | Sweet Bocas Trench 5 | -82.272318 | 9.359966667 | URS | Acropora | Acropora cervicornis | 444.03 |
| MC_9 | AT13-6-2- | Holocene | Sweet Bocas Trench 5 | -82.272318 | 9.359966667 | URS | Unknown | unknown | 458.95 |
| MC_9 | AT13-6-2- | Holocene | Sweet Bocas Trench 5 | -82.272318 | 9.359966667 | URS | Porites | Porites spp_dfp | 309.63 |
| MC_9 | AT13-6-2- | Holocene | Sweet Bocas Trench 5 | -82.272318 | 9.359966667 | URS | Orbicella | Orbicella spp | 1701.60 |

| | | | | | | | | | |
|---|---|---|---|---|---|---|---|---|---|
| NM_14 | AT13-6-3- | Holocene | Sweet Bocas Trench 5 | -82.272318 | 9.359966667 | URS | Oculina | Oculina spp | 0.15 |
| NM_14 | AT13-6-3- | Holocene | Sweet Bocas Trench 5 | -82.272318 | 9.359966667 | URS | cup | cup coral | 0.48 |
| NM_14 | AT13-6-3- | Holocene | Sweet Bocas Trench 5 | -82.272318 | 9.359966667 | URS | Cladocora | Cladocora arbuscula | 1.23 |
| NM_14 | AT13-6-3- | Holocene | Sweet Bocas Trench 5 | -82.272318 | 9.359966667 | URS | Millepora | Millepora spp | 12.14 |
| NM_14 | AT13-6-3- | Holocene | Sweet Bocas Trench 5 | -82.272318 | 9.359966667 | URS | Favia | Favia fragum | 15.21 |
| NM_14 | AT13-6-3- | Holocene | Sweet Bocas Trench 5 | -82.272318 | 9.359966667 | URS | Manicina | Manicina areolata | 50.46 |
| NM_14 | AT13-6-3- | Holocene | Sweet Bocas Trench 5 | -82.272318 | 9.359966667 | URS | Porites | Porites colonensis | 62.12 |
| NM_14 | AT13-6-3- | Holocene | Sweet Bocas Trench 5 | -82.272318 | 9.359966667 | URS | Agaricia | Agaricia spp | 118.87 |
| NM_14 | AT13-6-3- | Holocene | Sweet Bocas Trench 5 | -82.272318 | 9.359966667 | URS | Orbicella | Orbicella spp | 206.44 |
| NM_14 | AT13-6-3- | Holocene | Sweet Bocas Trench 5 | -82.272318 | 9.359966667 | URS | Unknown | unknown | 275.95 |
| NM_14 | AT13-6-3- | Holocene | Sweet Bocas Trench 5 | -82.272318 | 9.359966667 | URS | | other not coral | 470.10 |
| NM_14 | AT13-6-3- | Holocene | Sweet Bocas Trench 5 | -82.272318 | 9.359966667 | URS | Porites | Porites spp_dfp | 836.72 |
| NM_14 | AT13-6-3- | Holocene | Sweet Bocas Trench 5 | -82.272318 | 9.359966667 | URS | Acropora | Acropora cervicornis | 947.10 |
| MC_10 | AT13-6-6- | Holocene | Sweet Bocas Trench 5 | -82.272318 | 9.359966667 | URS | Madracis | Madracis spp | 0.18 |
| MC_10 | AT13-6-6- | Holocene | Sweet Bocas Trench 5 | -82.272318 | 9.359966667 | URS | Favia | Favia fragum | 0.94 |
| MC_10 | AT13-6-6- | Holocene | Sweet Bocas Trench 5 | -82.272318 | 9.359966667 | URS | | other not coral | 2.94 |
| MC_10 | AT13-6-6- | Holocene | Sweet Bocas Trench 5 | -82.272318 | 9.359966667 | URS | Cladocora | Cladocora arbuscula | 3.38 |
| MC_10 | AT13-6-6- | Holocene | Sweet Bocas Trench 5 | -82.272318 | 9.359966667 | URS | Millepora | Millepora spp | 6.52 |
| MC_10 | AT13-6-6- | Holocene | Sweet Bocas Trench 5 | -82.272318 | 9.359966667 | URS | Orbicella | Orbicella spp | 10.55 |
| MC_10 | AT13-6-6- | Holocene | Sweet Bocas Trench 5 | -82.272318 | 9.359966667 | URS | Agaricia | Agaricia spp | 37.01 |
| MC_10 | AT13-6-6- | Holocene | Sweet Bocas Trench 5 | -82.272318 | 9.359966667 | URS | Acropora | Acropora cervicornis | 109.23 |
| MC_10 | AT13-6-6- | Holocene | Sweet Bocas Trench 5 | -82.272318 | 9.359966667 | URS | Unknown | unknown | 256.61 |
| MC_10 | AT13-6-6- | Holocene | Sweet Bocas Trench 5 | -82.272318 | 9.359966667 | URS | Porites | Porites spp_dfp | 1269.00 |
| MC_11 | AT13-8-1- | Holocene | Sweet Bocas Trench 7 | -82.27273333 | 9.359683333 | URS | cup | cup coral | 0.00 |
| MC_11 | AT13-8-1- | Holocene | Sweet Bocas Trench 7 | -82.27273333 | 9.359683333 | URS | Leptoseris | Leptoseris cucullata | 0.71 |
| MC_11 | AT13-8-1- | Holocene | Sweet Bocas Trench 7 | -82.27273333 | 9.359683333 | URS | | other not coral | 0.82 |
| MC_11 | AT13-8-1- | Holocene | Sweet Bocas Trench 7 | -82.27273333 | 9.359683333 | URS | Oculina | Oculina diffusa | 0.828 |
| MC_11 | AT13-8-1- | Holocene | Sweet Bocas Trench 7 | -82.27273333 | 9.359683333 | URS | Cladocora | Cladocora arbuscula | 1.98 |
| MC_11 | AT13-8-1- | Holocene | Sweet Bocas Trench 7 | -82.27273333 | 9.359683333 | URS | Madracis | Madracis spp | 2.03 |
| MC_11 | AT13-8-1- | Holocene | Sweet Bocas Trench 7 | -82.27273333 | 9.359683333 | URS | Favia | Favia fragum | 4.05 |
| MC_11 | AT13-8-1- | Holocene | Sweet Bocas Trench 7 | -82.27273333 | 9.359683333 | URS | Millepora | Millepora spp | 4.22 |
| MC_11 | AT13-8-1- | Holocene | Sweet Bocas Trench 7 | -82.27273333 | 9.359683333 | URS | Unknown | unknown | 119.65 |
| MC_11 | AT13-8-1- | Holocene | Sweet Bocas Trench 7 | -82.27273333 | 9.359683333 | URS | Acropora | Acropora cervicornis | 182.83 |
| MC_11 | AT13-8-1- | Holocene | Sweet Bocas Trench 7 | -82.27273333 | 9.359683333 | URS | Orbicella | Orbicella spp | 497.31 |
| MC_11 | AT13-8-1- | Holocene | Sweet Bocas Trench 7 | -82.27273333 | 9.359683333 | URS | Porites | Porites spp_dfp | 516.08 |
| MC_11 | AT13-8-1- | Holocene | Sweet Bocas Trench 7 | -82.27273333 | 9.359683333 | URS | Agaricia | Agaricia spp | 1227.63 |
| MC_14 | AT13-8-2- | Holocene | Sweet Bocas Trench 7 | -82.27273333 | 9.359683333 | URS | Madracis | Madracis spp | 1.42 |
| MC_14 | AT13-8-2- | Holocene | Sweet Bocas Trench 7 | -82.27273333 | 9.359683333 | URS | Cladocora | Cladocora arbuscula | 2.60 |
| MC_14 | AT13-8-2- | Holocene | Sweet Bocas Trench 7 | -82.27273333 | 9.359683333 | URS | Porites | Porites astreoides | 3.49 |
| MC_14 | AT13-8-2- | Holocene | Sweet Bocas Trench 7 | -82.27273333 | 9.359683333 | URS | Leptoseris | Leptoseris cucullata | 3.96 |
| MC_14 | AT13-8-2- | Holocene | Sweet Bocas Trench 7 | -82.27273333 | 9.359683333 | URS | | other not coral | 5.62 |
| MC_14 | AT13-8-2- | Holocene | Sweet Bocas Trench 7 | -82.27273333 | 9.359683333 | URS | Millepora | Millepora spp | 6.38 |
| MC_14 | AT13-8-2- | Holocene | Sweet Bocas Trench 7 | -82.27273333 | 9.359683333 | URS | unknown | unknown | 88.02 |
| MC_14 | AT13-8-2- | Holocene | Sweet Bocas Trench 7 | -82.27273333 | 9.359683333 | URS | Manicina | Manicina areolata | 142.29 |
| MC_14 | AT13-8-2- | Holocene | Sweet Bocas Trench 7 | -82.27273333 | 9.359683333 | URS | Porites | Porites spp_dfp | 247.52 |
| MC_14 | AT13-8-2- | Holocene | Sweet Bocas Trench 7 | -82.27273333 | 9.359683333 | URS | Orbicella | Orbicella spp | 320.26 |
| MC_14 | AT13-8-2- | Holocene | Sweet Bocas Trench 7 | -82.27273333 | 9.359683333 | URS | Acropora | Acropora cervicornis | 468.12 |
| MC_14 | AT13-8-2- | Holocene | Sweet Bocas Trench 7 | -82.27273333 | 9.359683333 | URS | Agaricia | Agaricia spp | 781.48 |
| MC_19 | AT13-9-1- | Holocene | Sweet Bocas Trench 8 | -82.27365 | 9.360183333 | URS | Leptoseris | Leptoseris cucullata | 0.39 |
| MC_19 | AT13-9-1- | Holocene | Sweet Bocas Trench 8 | -82.27365 | 9.360183333 | URS | Millepora | Millepora spp | 1.04 |
| MC_19 | AT13-9-1- | Holocene | Sweet Bocas Trench 8 | -82.27365 | 9.360183333 | URS | Favia | Favia fragum | 1.06 |
| MC_19 | AT13-9-1- | Holocene | Sweet Bocas Trench 8 | -82.27365 | 9.360183333 | URS | Manicina | Manicina areolata | 1.24 |
| MC_19 | AT13-9-1- | Holocene | Sweet Bocas Trench 8 | -82.27365 | 9.360183333 | URS | Cladocora | Cladocora arbuscula | 1.41 |
| MC_19 | AT13-9-1- | Holocene | Sweet Bocas Trench 8 | -82.27365 | 9.360183333 | URS | | other not coral | 5.00 |
| MC_19 | AT13-9-1- | Holocene | Sweet Bocas Trench 8 | -82.27365 | 9.360183333 | URS | Eusmilia | Eusmilia fastigiata | 6.80 |
| MC_19 | AT13-9-1- | Holocene | Sweet Bocas Trench 8 | -82.27365 | 9.360183333 | URS | Unknown | unknown | 97.90 |
| MC_19 | AT13-9-1- | Holocene | Sweet Bocas Trench 8 | -82.27365 | 9.360183333 | URS | Agaricia | Agaricia spp | 225.71 |

| | | | | | | | | | |
|---|---|---|---|---|---|---|---|---|---|
| MC_19 | AT13-9-1- | Holocene | Sweet Bocas Trench 8 | -82.27365 | 9.360183333 | URS | Porites | Porites spp_dfp | 1621.63 |
| MC_18 | AT13-9-2- | Holocene | Sweet Bocas Trench 8 | -82.27365 | 9.360183333 | URS | cup | cup coral | 0.05 |
| MC_18 | AT13-9-2- | Holocene | Sweet Bocas Trench 8 | -82.27365 | 9.360183333 | URS | Favia | Favia fragum | 0.52 |
| MC_18 | AT13-9-2- | Holocene | Sweet Bocas Trench 8 | -82.27365 | 9.360183333 | URS | Cladocora | Cladocora arbuscula | 2.26 |
| MC_18 | AT13-9-2- | Holocene | Sweet Bocas Trench 8 | -82.27365 | 9.360183333 | URS | Leptoseris | Leptoseris cucullata | 5.30 |
| MC_18 | AT13-9-2- | Holocene | Sweet Bocas Trench 8 | -82.27365 | 9.360183333 | URS | Manicina | Manicina areolata | 5.98 |
| MC_18 | AT13-9-2- | Holocene | Sweet Bocas Trench 8 | -82.27365 | 9.360183333 | URS | | other not coral | 11.02 |
| MC_18 | AT13-9-2- | Holocene | Sweet Bocas Trench 8 | -82.27365 | 9.360183333 | URS | Unknown | unknown | 119.32 |
| MC_18 | AT13-9-2- | Holocene | Sweet Bocas Trench 8 | -82.27365 | 9.360183333 | URS | Agaricia | Agaricia spp | 221.59 |
| MC_18 | AT13-9-2- | Holocene | Sweet Bocas Trench 8 | -82.27365 | 9.360183333 | URS | Porites | Porites spp_dfp | 1455.73 |
| ML_11 | AT14-1-1- | Sub Recent | Casa Blanca new | -82.27989 | 9.36194 | 2.4 | Madracis | Madracis asperula | 6.54 |
| ML_11 | AT14-1-1- | Sub Recent | Casa Blanca new | -82.27989 | 9.36194 | 2.4 | Leptoseris | Leptoseris cucullata | 0.28 |
| ML_11 | AT14-1-1- | Sub Recent | Casa Blanca new | -82.27989 | 9.36194 | 2.4 | Agaricia | Agaricia spp | 7.75 |
| ML_11 | AT14-1-1- | Sub Recent | Casa Blanca new | -82.27989 | 9.36194 | 2.4 | | other not coral | 124.00 |
| ML_11 | AT14-1-1- | Sub Recent | Casa Blanca new | -82.27989 | 9.36194 | 2.4 | Acropora | Acropora cervicornis | 125.00 |
| ML_11 | AT14-1-1- | Sub Recent | Casa Blanca new | -82.27989 | 9.36194 | 2.4 | Porites | Porites spp_dfp | 417.04 |
| ML_11 | AT14-1-1- | Sub Recent | Casa Blanca new | -82.27989 | 9.36194 | 2.4 | Unknown | unknown | 685.90 |
| NM_7 | AT14-1-2- | Sub Recent | Casa Blanca new | -82.27989 | 9.36194 | 3.1 | Millepora | Millepora spp | 0.46 |
| NM_7 | AT14-1-2- | Sub Recent | Casa Blanca new | -82.27989 | 9.36194 | 3.1 | Madracis | Madracis decactis | 0.68 |
| NM_7 | AT14-1-2- | Sub Recent | Casa Blanca new | -82.27989 | 9.36194 | 3.1 | Agaricia | Agaricia spp | 7.56 |
| NM_7 | AT14-1-2- | Sub Recent | Casa Blanca new | -82.27989 | 9.36194 | 3.1 | Unknown | unknown | 27.42 |
| NM_7 | AT14-1-2- | Sub Recent | Casa Blanca new | -82.27989 | 9.36194 | 3.1 | Acropora | Acropora cervicornis | 89.23 |
| NM_7 | AT14-1-2- | Sub Recent | Casa Blanca new | -82.27989 | 9.36194 | 3.1 | Porites | Porites spp_dfp | 92.41 |
| NM_7 | AT14-1-2- | Sub Recent | Casa Blanca new | -82.27989 | 9.36194 | 3.1 | | other not coral | 318.73 |
| NM_8 | AT14-1-3- | Sub Recent | Casa Blanca new | -82.27989 | 9.36194 | 2.6 | Madracis | Madracis asperula | 2.05 |
| NM_8 | AT14-1-3- | Sub Recent | Casa Blanca new | -82.27989 | 9.36194 | 2.6 | Favia | Favia fragum | 0.88 |
| NM_8 | AT14-1-3- | Sub Recent | Casa Blanca new | -82.27989 | 9.36194 | 2.6 | Millepora | Millepora spp | 1.49 |
| NM_8 | AT14-1-3- | Sub Recent | Casa Blanca new | -82.27989 | 9.36194 | 2.6 | Porites | Porites colonensis | 2.04 |
| NM_8 | AT14-1-3- | Sub Recent | Casa Blanca new | -82.27989 | 9.36194 | 2.6 | Eusmilia | Eusmilia fastigiata | 5.06 |
| NM_8 | AT14-1-3- | Sub Recent | Casa Blanca new | -82.27989 | 9.36194 | 2.6 | Agaricia | Agaricia spp | 18.63 |
| NM_8 | AT14-1-3- | Sub Recent | Casa Blanca new | -82.27989 | 9.36194 | 2.6 | Acropora | Acropora cervicornis | 83.08 |
| NM_8 | AT14-1-3- | Sub Recent | Casa Blanca new | -82.27989 | 9.36194 | 2.6 | Unknown | unknown | 195.45 |
| NM_8 | AT14-1-3- | Sub Recent | Casa Blanca new | -82.27989 | 9.36194 | 2.6 | | other not coral | 329.16 |
| NM_8 | AT14-1-3- | Sub Recent | Casa Blanca new | -82.27989 | 9.36194 | 2.6 | Porites | Porites spp_dfp | 999.06 |
| NM_9_MC_2 | AT14-1-7- | Sub Recent | Casa Blanca new | -82.27989 | 9.36194 | 3.9 | Montastraea | Montastraea annularis | 0.18 |
| NM_9_MC_2 | AT14-1-7- | Sub Recent | Casa Blanca new | -82.27989 | 9.36194 | 3.9 | Oculina | Oculina spp | 0.26 |
| NM_9_MC_2 | AT14-1-7- | Sub Recent | Casa Blanca new | -82.27989 | 9.36194 | 3.9 | Eusmilia | Eusmilia fastigiata | 0.48 |
| NM_9_MC_2 | AT14-1-7- | Sub Recent | Casa Blanca new | -82.27989 | 9.36194 | 3.9 | Leptoseris | Leptoseris cucullata | 1.14 |
| NM_9_MC_2 | AT14-1-7- | Sub Recent | Casa Blanca new | -82.27989 | 9.36194 | 3.9 | Favia | Favia fragum | 6.29 |
| NM_9_MC_2 | AT14-1-7- | Sub Recent | Casa Blanca new | -82.27989 | 9.36194 | 3.9 | Madracis | Madracis spp | 10.42 |
| NM_9_MC_2 | AT14-1-7- | Sub Recent | Casa Blanca new | -82.27989 | 9.36194 | 3.9 | Madracis | Madracis asperula | 15.66 |
| NM_9_MC_2 | AT14-1-7- | Sub Recent | Casa Blanca new | -82.27989 | 9.36194 | 3.9 | Acropora | Acropora cervicornis | 74.18 |
| NM_9_MC_2 | AT14-1-7- | Sub Recent | Casa Blanca new | -82.27989 | 9.36194 | 3.9 | Agaricia | Agaricia spp | 95.23 |
| NM_9_MC_2 | AT14-1-7- | Sub Recent | Casa Blanca new | -82.27989 | 9.36194 | 3.9 | Unknown | unknown | 482.48 |
| NM_9_MC_2 | AT14-1-7- | Sub Recent | Casa Blanca new | -82.27989 | 9.36194 | 3.9 | Porites | Porites spp_dfp | 618.86 |
| NM_9_MC_2 | AT14-1-7- | Sub Recent | Casa Blanca new | -82.27989 | 9.36194 | 3.9 | | other not coral | 668.30 |
| MC_3 | AT14-1-8- | Sub Recent | Casa Blanca new | -82.27989 | 9.36194 | 3.4 | Leptoseris | Leptoseris cucullata | 0.76 |
| MC_3 | AT14-1-8- | Sub Recent | Casa Blanca new | -82.27989 | 9.36194 | 3.4 | Madracis | Madracis spp | 3.35 |
| MC_3 | AT14-1-8- | Sub Recent | Casa Blanca new | -82.27989 | 9.36194 | 3.4 | Acropora | Acropora cervicornis | 7.17 |
| MC_3 | AT14-1-8- | Sub Recent | Casa Blanca new | -82.27989 | 9.36194 | 3.4 | Agaricia | Agaricia spp | 10.78 |
| MC_3 | AT14-1-8- | Sub Recent | Casa Blanca new | -82.27989 | 9.36194 | 3.4 | | other not coral | 250.10 |
| MC_3 | AT14-1-8- | Sub Recent | Casa Blanca new | -82.27989 | 9.36194 | 3.4 | Porites | Porites spp_dfp | 477.98 |
| MC_3 | AT14-1-8- | Sub Recent | Casa Blanca new | -82.27989 | 9.36194 | 3.4 | Unknown | unknown | 548.95 |
| NM_10_MC_5 | AT14-4-2- | Sub Recent | Airport Point | -82.25611 | 9.33611 | 5.2 | Cladocora | Cladocora arbuscula | 0.29 |
| NM_10_MC_5 | AT14-4-2- | Sub Recent | Airport Point | -82.25611 | 9.33611 | 5.2 | Eusmilia | Eusmilia fastigiata | 4.91 |
| NM_10_MC_5 | AT14-4-2- | Sub Recent | Airport Point | -82.25611 | 9.33611 | 5.2 | Millepora | Millepora spp | 0.13 |
| NM_10_MC_5 | AT14-4-2- | Sub Recent | Airport Point | -82.25611 | 9.33611 | 5.2 | cup | cup coral | 0.86 |

| | | | | | | | | | |
|---|---|---|---|---|---|---|---|---|---|
| NM_10_MC_5 | AT14-4-2- | Sub Recent | Airport Point | -82.25611 | 9.33611 | 5.2 | Porites | Porites colonensis | 1.01 |
| NM_10_MC_5 | AT14-4-2- | Sub Recent | Airport Point | -82.25611 | 9.33611 | 5.2 | Leptoseris | Leptoseris cucullata | 1.09 |
| NM_10_MC_5 | AT14-4-2- | Sub Recent | Airport Point | -82.25611 | 9.33611 | 5.2 | Manicina | Manicina areolata | 1.57 |
| NM_10_MC_5 | AT14-4-2- | Sub Recent | Airport Point | -82.25611 | 9.33611 | 5.2 | Madracis | Madracis spp | 2.27 |
| NM_10_MC_5 | AT14-4-2- | Sub Recent | Airport Point | -82.25611 | 9.33611 | 5.2 | Favia | Favia fragum | 3.48 |
| NM_10_MC_5 | AT14-4-2- | Sub Recent | Airport Point | -82.25611 | 9.33611 | 5.2 | Oculina | Oculina spp | 3.73 |
| NM_10_MC_5 | AT14-4-2- | Sub Recent | Airport Point | -82.25611 | 9.33611 | 5.2 | Agaricia | Agaricia spp | 437.26 |
| NM_10_MC_5 | AT14-4-2- | Sub Recent | Airport Point | -82.25611 | 9.33611 | 5.2 | | other not coral | 551.30 |
| NM_10_MC_5 | AT14-4-2- | Sub Recent | Airport Point | -82.25611 | 9.33611 | 5.2 | Unknown | unknown | 606.39 |
| NM_10_MC_5 | AT14-4-2- | Sub Recent | Airport Point | -82.25611 | 9.33611 | 5.2 | Porites | Porites spp_dfp | 756.28 |
| NM_11_MC_6 | AT14-4-3- | Sub Recent | Airport Point | -82.25611 | 9.33611 | 5.5 | Millepora | Millepora spp | 1.25 |
| NM_11_MC_6 | AT14-4-3- | Sub Recent | Airport Point | -82.25611 | 9.33611 | 5.5 | Porites | Porites colonensis | 1.29 |
| NM_11_MC_6 | AT14-4-3- | Sub Recent | Airport Point | -82.25611 | 9.33611 | 5.5 | Oculina | Oculina spp | 1.60 |
| NM_11_MC_6 | AT14-4-3- | Sub Recent | Airport Point | -82.25611 | 9.33611 | 5.5 | Leptoseris | Leptoseris cucullata | 1.75 |
| NM_11_MC_6 | AT14-4-3- | Sub Recent | Airport Point | -82.25611 | 9.33611 | 5.5 | cup | cup coral | 2.20 |
| NM_11_MC_6 | AT14-4-3- | Sub Recent | Airport Point | -82.25611 | 9.33611 | 5.5 | Favia | Favia fragum | 3.02 |
| NM_11_MC_6 | AT14-4-3- | Sub Recent | Airport Point | -82.25611 | 9.33611 | 5.5 | Eusmilia | Eusmilia fastigiata | 3.19 |
| NM_11_MC_6 | AT14-4-3- | Sub Recent | Airport Point | -82.25611 | 9.33611 | 5.5 | Madracis | Madracis spp | 5.15 |
| NM_11_MC_6 | AT14-4-3- | Sub Recent | Airport Point | -82.25611 | 9.33611 | 5.5 | Agaricia | Agaricia spp | 259.50 |
| NM_11_MC_6 | AT14-4-3- | Sub Recent | Airport Point | -82.25611 | 9.33611 | 5.5 | Unknown | unknown | 553.38 |
| NM_11_MC_6 | AT14-4-3- | Sub Recent | Airport Point | -82.25611 | 9.33611 | 5.5 | Porites | Porites spp_dfp | 688.46 |
| NM_11_MC_6 | AT14-4-3- | Sub Recent | Airport Point | -82.25611 | 9.33611 | 5.5 | | other not coral | 721.30 |
| MC_7 | AT14-4-4- | Sub Recent | Airport Point | -82.25611 | 9.33611 | 5.4 | Cladocora | Cladocora arbuscula | 0.19 |
| MC_7 | AT14-4-4- | Sub Recent | Airport Point | -82.25611 | 9.33611 | 5.4 | cup | cup coral | 1.72 |
| MC_7 | AT14-4-4- | Sub Recent | Airport Point | -82.25611 | 9.33611 | 5.4 | Oculina | Oculina diffusa | 1.87 |
| MC_7 | AT14-4-4- | Sub Recent | Airport Point | -82.25611 | 9.33611 | 5.4 | Leptoseris | Leptoseris cucullata | 2.871 |
| MC_7 | AT14-4-4- | Sub Recent | Airport Point | -82.25611 | 9.33611 | 5.4 | Madracis | Madracis spp | 4.64 |
| MC_7 | AT14-4-4- | Sub Recent | Airport Point | -82.25611 | 9.33611 | 5.4 | Favia | Favia fragum | 5.88 |
| MC_7 | AT14-4-4- | Sub Recent | Airport Point | -82.25611 | 9.33611 | 5.4 | Manicina | Manicina areolata | 8.18 |
| MC_7 | AT14-4-4- | Sub Recent | Airport Point | -82.25611 | 9.33611 | 5.4 | Millepora | Millepora spp | 9.73 |
| MC_7 | AT14-4-4- | Sub Recent | Airport Point | -82.25611 | 9.33611 | 5.4 | Porites | Porites astreoides | 59.79 |
| MC_7 | AT14-4-4- | Sub Recent | Airport Point | -82.25611 | 9.33611 | 5.4 | Agaricia | Agaricia spp | 133.09 |
| MC_7 | AT14-4-4- | Sub Recent | Airport Point | -82.25611 | 9.33611 | 5.4 | Porites | Porites spp_dfp | 787.39 |
| MC_7 | AT14-4-4- | Sub Recent | Airport Point | -82.25611 | 9.33611 | 5.4 | | other not coral | 796.49 |
| MC_7 | AT14-4-4- | Sub Recent | Airport Point | -82.25611 | 9.33611 | 5.4 | Unknown | unknown | 889.51 |
| NM_13 | AT14-4-5- | Sub Recent | Airport Point | -82.25611 | 9.33611 | 5.1 | cup | cup coral | 0.12 |
| NM_13 | AT14-4-5- | Sub Recent | Airport Point | -82.25611 | 9.33611 | 5.1 | Madracis | Madracis spp | 0.16 |
| NM_13 | AT14-4-5- | Sub Recent | Airport Point | -82.25611 | 9.33611 | 5.1 | Cladocora | Cladocora arbuscula | 0.35 |
| NM_13 | AT14-4-5- | Sub Recent | Airport Point | -82.25611 | 9.33611 | 5.1 | Leptoseris | Leptoseris cucullata | 0.95 |
| NM_13 | AT14-4-5- | Sub Recent | Airport Point | -82.25611 | 9.33611 | 5.1 | Scolymia | Scolymia cubensis | 1.27 |
| NM_13 | AT14-4-5- | Sub Recent | Airport Point | -82.25611 | 9.33611 | 5.1 | Porites | Porites colonensis | 1.46 |
| NM_13 | AT14-4-5- | Sub Recent | Airport Point | -82.25611 | 9.33611 | 5.1 | Favia | Favia fragum | 7.82 |
| NM_13 | AT14-4-5- | Sub Recent | Airport Point | -82.25611 | 9.33611 | 5.1 | Eusmilia | Eusmilia fastigiata | 28.32 |
| NM_13 | AT14-4-5- | Sub Recent | Airport Point | -82.25611 | 9.33611 | 5.1 | Oculina | Oculina spp | 35.02 |
| NM_13 | AT14-4-5- | Sub Recent | Airport Point | -82.25611 | 9.33611 | 5.1 | Manicina | Manicina areolata | 87.14 |
| NM_13 | AT14-4-5- | Sub Recent | Airport Point | -82.25611 | 9.33611 | 5.1 | Agaricia | Agaricia spp | 613.5365 |
| NM_13 | AT14-4-5- | Sub Recent | Airport Point | -82.25611 | 9.33611 | 5.1 | | other not coral | 758.90 |
| NM_13 | AT14-4-5- | Sub Recent | Airport Point | -82.25611 | 9.33611 | 5.1 | Unknown | unknown | 914.58 |
| NM_13 | AT14-4-5- | Sub Recent | Airport Point | -82.25611 | 9.33611 | 5.1 | Porites | Porites spp_dfp | 1325.18 |
| ML_12_MC_1 | AT14-4-6- | Sub Recent | Airport Point | -82.25611 | 9.33611 | 5 | Leptoseris | Leptoseris cucullata | 0.74 |
| ML_12_MC_1 | AT14-4-6- | Sub Recent | Airport Point | -82.25611 | 9.33611 | 5 | Cladocora | Cladocora arbuscula | 0.52 |
| ML_12_MC_1 | AT14-4-6- | Sub Recent | Airport Point | -82.25611 | 9.33611 | 5 | cup | cup coral | 0.14 |
| ML_12_MC_1 | AT14-4-6- | Sub Recent | Airport Point | -82.25611 | 9.33611 | 5 | Favia | Favia fragum | 2.62 |
| ML_12_MC_1 | AT14-4-6- | Sub Recent | Airport Point | -82.25611 | 9.33611 | 5 | Manicina | Manicina areolata | 2.87 |
| ML_12_MC_1 | AT14-4-6- | Sub Recent | Airport Point | -82.25611 | 9.33611 | 5 | Madracis | Madracis spp | 5.86 |
| ML_12_MC_1 | AT14-4-6- | Sub Recent | Airport Point | -82.25611 | 9.33611 | 5 | Oculina | Oculina spp | 5.86 |
| ML_12_MC_1 | AT14-4-6- | Sub Recent | Airport Point | -82.25611 | 9.33611 | 5 | Eusmilia | Eusmilia fastigiata | 7.98 |

| | | | | | | | | | |
|---|---|---|---|---|---|---|---|---|---|
| ML_12_MC_1 | AT14-4-6- | Sub Recent | Airport Point | -82.25611 | 9.33611 | 5 | Millepora | Millepora spp | 15.91 |
| ML_12_MC_1 | AT14-4-6- | Sub Recent | Airport Point | -82.25611 | 9.33611 | 5 | Agaricia | Agaricia spp | 341.07 |
| ML_12_MC_1 | AT14-4-6- | Sub Recent | Airport Point | -82.25611 | 9.33611 | 5 | Porites | Porites spp_dfp | 489.92 |
| ML_12_MC_1 | AT14-4-6- | Sub Recent | Airport Point | -82.25611 | 9.33611 | 5 | | other not coral | 979.60 |
| ML_12_MC_1 | AT14-4-6- | Sub Recent | Airport Point | -82.25611 | 9.33611 | 5 | Unknown | unknown | 1523.13 |
